# Decreased Directed Functional Connectivity in the Psychedelic State

**DOI:** 10.1101/703660

**Authors:** Lionel Barnett, Suresh D. Muthukumaraswamy, Robin L. Carhart-Harris, Anil K. Seth

## Abstract

Neuroimaging studies of the psychedelic state offer a unique window onto the neural basis of conscious perception and selfhood. Despite well understood pharmacological mechanisms of action, the large-scale changes in neural dynamics induced by psychedelic compounds remain poorly understood. Using source-localised, steady-state MEG recordings, we describe changes in functional connectivity following the controlled administration of LSD, psilocybin and low-dose ketamine, as well as, for comparison, the (non-psychedelic) anticonvulsant drug tiagabine. We compare both undirected and directed measures of functional connectivity between placebo and drug conditions. We observe a general decrease in directed functional connectivity for all three psychedelics, as measured by Granger causality, throughout the brain. These data support the view that the psychedelic state involves a breakdown in patterns of functional organisation or information flow in the brain. In the case of LSD, the decrease in directed functional connectivity is coupled with an increase in undirected functional connectivity, which we measure using correlation and coherence. This surprising opposite movement of directed and undirected measures is of more general interest for functional connectivity analyses, which we interpret using analytical modelling. Overall, our results uncover the neural dynamics of information flow in the psychedelic state, and highlight the importance of comparing multiple measures of functional connectivity when analysing time-resolved neuroimaging data.

## 1 Introduction

Psychedelic drugs impact profoundly on conscious experiences of world and self, providing a unique opportunity to examine the neural and cognitive bases of these subjective phenomena. The classical psychedelics, notably LSD and psilocybin, have well-characterised pharmacological mechanisms of action. Both are serotonin (5HT2AR) agonists, while ketamine - which at low (sub-anesthetic) doses also has psychedelic-like properties - is primarily an NMDA antagonist (Sleigh et al., 2014). Many studies now link the subjective effects of psychedelics to altered activity at these receptors (Preller et al., 2018; Deco et al., 2018). However, despite these links, the large-scale changes in neural dynamics that underlie the dramatic subjective effects of psychedelics remain poorly understood.

One window onto large-scale neural dynamics is to measure functional connectivity (FC). FC between brain regions refers to the existence of statistical dependencies in their activity (Seth et al., 2015). Several previous FC analyses of the psychedelic state analyse resting-state fMRI data obtained from healthy volunteers under either psilocybin or LSD. These studies have revealed a reorganisation of connectivity within and between resting-state networks, generally showing decreased connectivity within resting-state networks and increased connectivity across such networks (Carhart-Harris et al., 2012; Tagliazucchi et al., 2016; Carhart-Harris et al., 2016; Müller et al., 2017, 2018; Kaelen et al., 2016), see also Preller et al. (2018). For example, decreased connectivity within the default-mode network has been associated with subjective ego dissolution (Smigielski et al., 2019), as has increased connectivity across normally functionally segregated networks (Tagliazucchi et al., 2016).

Despite the insights provided by these studies there are inherent limitations on FC analysis of fMRI data imposed by the poor temporal resolution of such data, induced both by low sampling frequencies and the slow dynamics of the hemodynamic response. More fine-grained FC analysis is possible when analysing high-time resolution electrophysiological data, as can be obtained using EEG or MEG. Importantly, because of the high temporal resolution (as compared with fMRI), such data are particularly suitable for measuring and comparing *undirected* and *directed* measures of FC. MEG, in particular, is highly suitable for such analysis since unlike EEG it is not subject to confounds arising through volume conduction in the skull.

The distinction between directed and undirected FC is critical in providing a comprehensive picture of statistical dependencies between regional brain activities. Undirected measures, like correlation and coherence, are symmetric by definition. These measures can be thought of as reflecting “shared information” between variables, with mutual information being the most general case. Directed measures, such as Granger causality (GC), are generally not symmetric; these measures reflect “information flow” between variables, with transfer entropy (Schreiber, 2000; Paluš et al., 2001) being the most general case (Barnett et al., 2009; Barnett and Bossomaier, 2013). Note that both are distinct from measures of “effective connectivity (EC)” [for example, dynamic causal modelling (DCM, Friston et al., 2013)] which aim to describe the minimal causal circuit capable of reproducing some observed dynamics [see Muthukumaraswamy et al. (2015a) for an application of DCM to brain dynamics under ketamine]. Functional connectivity metrics, whether directed or undirected, describe relationships between dynamical variables of a system, whereas effective connectivity uses dynamics to make inferences about the underlying mechanisms in a system (Seth et al., 2015; Barnett et al., 2009; Barnett and Seth, 2014; Friston et al., 2013).

A few studies have applied fine-grained FC analyses to neuroimaging data obtained in the investigations of the psychedelic state. In an EEG study, Kometer et al. (2015) report increased (undirected) phase synchronisation under psilocybin. Using MEG, Rivolta et al. (2015) find increased (directed) transfer entropy in thalamocortical networks under low-dose ketamine. Transfer entropy has also been applied to EEG data obtained under ayahuasca (another serotonergic psychedelic), revealing alterations in anterior-to-posterior and posterior-to-anterior information flow (Alonso et al., 2015). In an EEG study on the functional effects of sub-anesthetic ketamine, Vlisides et al. (2017) report (undirected) theta phase locking between anterior and posterior regions as measured by the weighted phase lag index, along with reduced anterior-to-posterior directed connectivity as measured by the directed phase lag index (Stam and van Straaten, 2012). However, a comprehensive picture of how directed and undirected FC change together, under a range of psychedelics, and compared against non-psychedelic controls, is still lacking.

We therefore set out to characterise the large-scale alterations in FC in the psychedelic state by analysing source-localised MEG data, using both directed and undirected FC measures on the same data, and comparing a range of psychedelics as well as a non-psychedelic control. These data had been previously obtained from healthy volunteers under controlled intravenous infusion of one of LSD, psilocybin, ketamine, or tiagabine (the control condition; tiagabine is a non-psychedelic GABA reuptake inhibitor, which is clinically employed as an anticonvulsant). MEG was carried out on healthy volunteers in a non-task resting state, and each volunteer underwent a placebo session as well as a drug session. We measured both undirected FC (correlation in the time domain, coherence in the frequency domain) and directed FC (Granger causality, in both time and frequency domains) on the same data. Our analyses were, where possible, fully conditional to control for indirect influences, which can affect inferences about FC in multivariate systems. Our primary question was whether the psychedelics would elicit reliable changes in directed and undirected FC between and within cortical regions, as compared with placebo, and as compared with the non-psychedelic drug (tiagabine) control. All analyses were treated as exploratory.

Anticipating results, we find a decrease in directed FC generally between brain regions, for all psychedelic compounds, but not for tiagabine (control). In some instances these decreases are accompanied by increases in undirected FC. We interpret these results as suggesting decreased neural information flow in the psychedelic state, consistent with perspectives that emphasise increasing disorder and functional disorganisation underlying psychedelic experience (Carhart-Harris et al., 2012; Carhart-Harris, 2018; Carhart-Harris and Friston, 2019; Schartner et al., 2017).

## 2 Materials and Methods

### 2.1 Experimental procedure, data acquisition and preprocessing

The MEG datasets for resting-state LSD, psilocybin (PSI), ketamine (KET) and tiagabine (TGB) used in this article have all previously been analysed in published studies. For an overview of the respective MEG acquisition and preprocessing procedures for the psychedelic compounds, see Schartner et al. (2017). Full details are given in Carhart-Harris et al. (2016) [LSD], Muthukumaraswamy et al. (2013) [PSI], Muthukumaraswamy et al. (2015b) [KET], and Nutt et al. (2015) [TGB]. The MEG data were source-localised to 90 cortical regions according to the standard Automated Anatomical Labelling (AAL) brain atlas (Tzourio-Mazoyer et al., 2002). For some analyses, the 90 AAL regions were grouped into the larger anatomical regions listed in Table 1. We refer to the AAL regions as “sources”, and the larger regions in Table 1 as “ROIs”.

**Table 1:** Large anatomical regions (ROIs).

| Label | Name | AAL sources |
| --- | --- | --- |
| Cin | Cingulate | 31-36 |
| Fro | Frontal | 3-16, 19-28 |
| Lim | Limbic | 37-42 |
| Occ | Occipital | 43-56 |
| Par | Parietal | 59-70 |
| Sen | Sensorimotor | 1-2,17-18,57-58 |
| Sub | Subcortical | 71-78 |
| Tem | Temporal | 29-30,79-90 |

All analyses are based on resting-state recordings post-infusion of drug and of placebo. The recording length varies by drug (7 − 14 min for LSD, 5 min for PSI and TGB, and 10 min for KET). Originally sampled at 600 Hz or 1200 Hz (depending on the study), the data was high-pass filtered at 1 Hz to suppress slow transients, low-pass filtered at 150 Hz, and downsampled to 300 Hz (Barnett and Seth, 2011)^1^. Line noise was suppressed by subtracting a least-squares-fit sinusoidal signal at 50 Hz and harmonics. The MEG time series were segmented into 2 s epochs comprising 600 Hz ×2s = 1200 observations, and artefacted epochs discarded (see above refs. for details); the number of usable epochs thus varies with drug, subject and condition (i.e., drug or placebo); numbers are given in Table 2. Epochs are analysed as stationary multi-trial (also known as “panel”) data: the assumption is that all 2 s epochs for a given drug/subject/condition are realisations of the same underlying stochastic process, and may therefore be pooled for statistical estimation. Although this assumption is difficult to validate rigorously, in the present case treating the data as multi-trial yielded stable autoregressive models (Section 2.2.2), indicating consistent statistical properties across epochs. It is also a reasonable assumption given the relatively short length of the MEG recordings relative to the duration of the subjective drug effects, which reliably extended long beyond the recording periods. With the exception of spectral power estimation, epochs were normalised per drug/subject/condition by the pooled mean and variance. (We note that, for multi-trial estimation, epochs should not be normalised individually by per-epoch mean and variance, as this is known to induce statistical bias.)

**Table 2:**
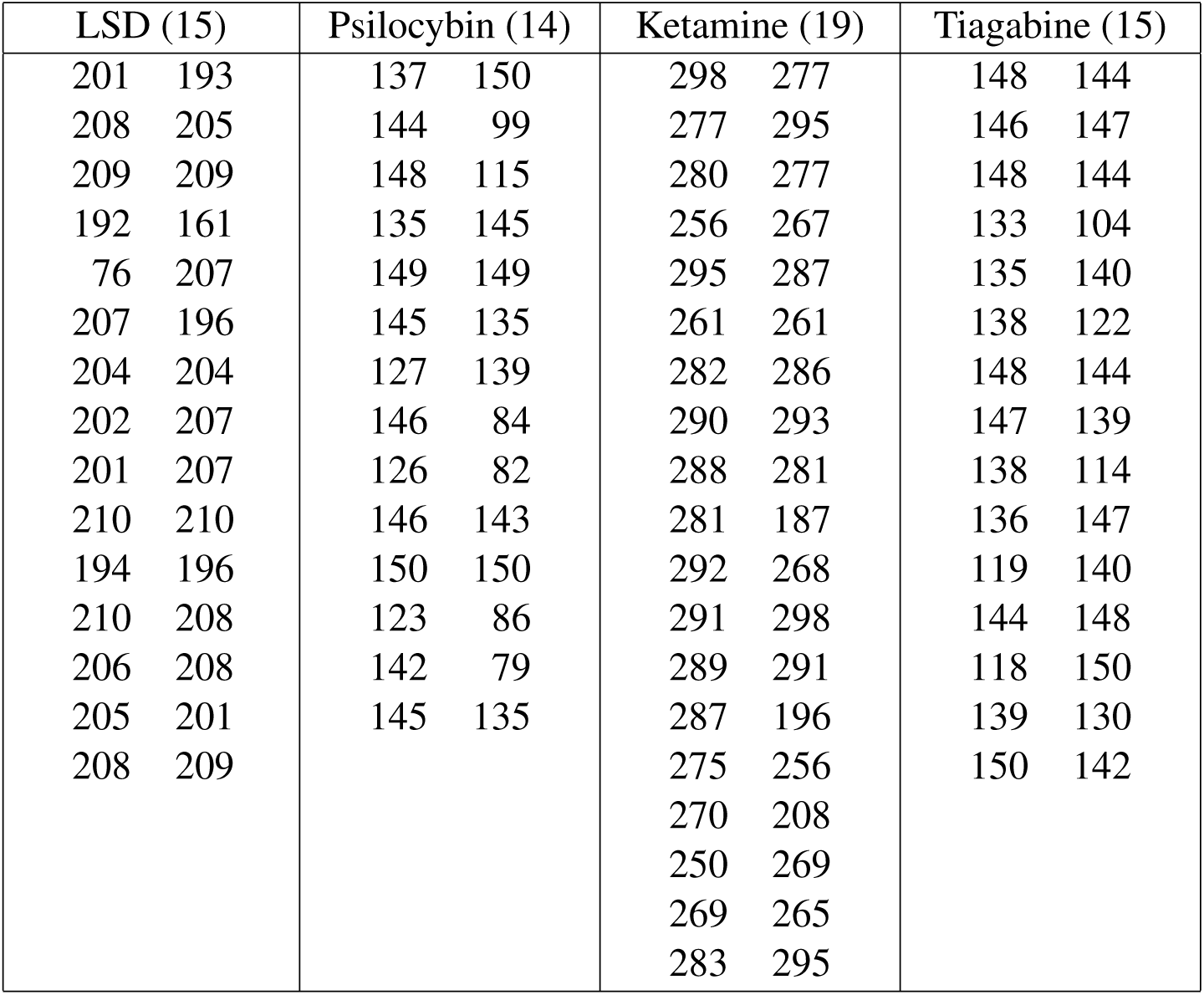
Number of epochs per drug by subject, after removal of artefacted epochs. Number in brackets is number of subjects per drug. Left-hand columns: placebo epochs per subject, right-hand columns: drug epochs per subject.

| LSD (15) | Psilocybin (14) | Ketamine (19) | Tiagabine (15) |
| --- | --- | --- | --- |
| 201 193 | 137 150 | 298 277 | 148 144 |
| 208 205 | 144 99 | 277 295 | 146 147 |
| 209 209 | 148 115 | 280 277 | 148 144 |
| 192 161 | 135 145 | 256 267 | 133 104 |
| 76 207 | 149 149 | 295 287 | 135 140 |
| 207 196 | 145 135 | 261 261 | 138 122 |
| 204 204 | 127 139 | 282 286 | 148 144 |
| 202 207 | 146 84 | 290 293 | 147 139 |
| 201 207 | 126 82 | 288 281 | 138 114 |
| 210 210 | 146 143 | 281 187 | 136 147 |
| 194 196 | 150 150 | 292 268 | 119 140 |
| 210 208 | 123 86 | 291 298 | 144 148 |
| 206 208 | 142 79 | 289 291 | 118 150 |
| 205 201 | 145 135 | 287 196 | 139 130 |
| 208 209 |  | 275 256 | 150 142 |
|  |  | 270 208 |  |
|  |  | 250 269 |  |
|  |  | 269 265 |  |
|  |  | 283 295 |  |

### 2.2 Functional connectivity analysis

We applied both directed and undirected measures of functional connectivity. In the time domain we measured correlation (undirected) and Granger causality (directed). In the frequency domain we measured coherence (undirected) and spectral Granger causality (directed). These statistics all have information-theoretic interpretations which are exact if the data are Gaussian (Barnett et al., 2009), and asymptotic otherwise (Barnett and Bossomaier, 2013). (The marginal distributions of the normalised MEG data are, in our case, approximately Gaussian.) Correlation equates to mutual information (Cover and Thomas, 1991), a measure of undirected shared information, while Granger causality equates to transfer entropy (Schreiber, 2000; Paluš et al., 2001), a measure of the rate of directed information flow between stochastic processes.

In multivariate situations, measures of FC can be confounded by indirect associations. That is, a variable *A* may appear to be correlated with (or to Granger-cause) a variable *B* when only *A* and *B* are measured, but may be revealed to be unrelated when a third (common influence) variable *C* is also included in the analysis. Therefore, where appropriate, we used conditional (or “partial”) FC measures, which control for indirect associations. (Note that it is not possible to control for indirect associations mediated by latent/unrecorded common influences, although their presence may potentially be inferred; *cf*. Section 4.) Conditioning in a highly multivariate context generally requires large amounts of data to achieve statistical power; this was unproblematic in the present analysis. Frequency-domain statistics are defined so that they may be integrated (averaged) across a given frequency band (Section 2.2.3) to yield measures of FC in specific frequency ranges (Table 3). For GC, the frequency-domain statistic integrates across the broadband spectrum to yield the corresponding time-domain statistic.

**Table 3:**
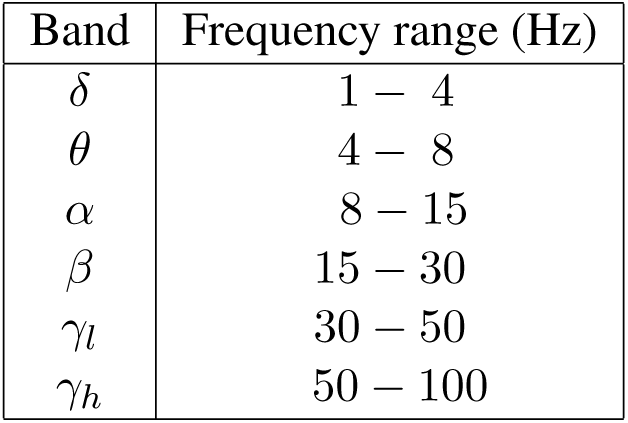
Standardised frequency bands. Frequencies above 100 Hz are omitted from *γ*_*h*_ as they are possibly compromised by roll-off from 100 Hz low-pass filter, and in any case were not considered functionally relevant.

For each drug, subject and condition we take each 2 s MEG epoch as a realisation of a stationary multivariate stochastic process *X*_*t*_ = {*X*_*it*_} where *t* indexes time steps and *i* indexes localised sources corresponding to the 90 AAL regions. The information-theoretic and frequency-domain measures described below are estimated under Gaussian assumptions as 2nd-order statistics, based on vector-autoregressive (VAR) modelling (Hamilton, 1994; Lütkepohl, 1993)^2^. Here, the (zero-mean) process *X*_*t*_ is modelled as a (finite-order, stable and invertible) vector autoregression

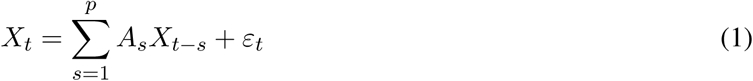

with white noise (iid, serially-uncorrelated) residuals *ε*_*t*_. The model parameters are the regression coefficients matrices *A*_1_, …, *A*_*p*_, and the variance-covariance matrix 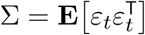. For each drug, subject and condition, a model order *p* must first be selected; here we used a likelihood-ratio F-test (Hamilton, 1994); *p* generally varied between 2 and 5 (Table 4). Model parameters were then identified from the normalised time-series data using an LWR maximum-likelihood estimator due to Morf et al. (1978).

**Table 4:** Model orders by drug per subject. Number in brackets is number of subjects per drug (Table 2). Left-hand columns: placebo, right-hand columns: drug.

| LSD (15) | Psilocybin (14) | Ketamine (19) | Tiagabine (15) |
| --- | --- | --- | --- |
| 3 3 | 4 4 | 4 3 | 4 5 |
| 4 3 | 4 3 | 3 2 | 4 4 |
| 3 3 | 4 3 | 4 3 | 4 4 |
| 3 3 | 4 4 | 3 3 | 4 5 |
| 4 3 | 4 4 | 4 3 | 5 4 |
| 3 3 | 3 3 | 3 3 | 4 4 |
| 3 3 | 3 3 | 4 3 | 4 4 |
| 3 3 | 4 4 | 4 3 | 4 4 |
| 4 3 | 4 3 | 3 2 | 4 5 |
| 3 3 | 4 4 | 4 3 | 5 4 |
| 4 3 | 4 4 | 3 3 | 4 4 |
| 4 4 | 4 4 | 4 3 | 4 4 |
| 4 3 | 3 3 | 3 3 | 4 4 |
| 3 3 | 4 4 | 4 3 | 5 4 |
| 3 3 |  | 4 3 | 4 4 |
|  |  | 3 3 |  |
|  |  | 3 3 |  |
|  |  | 3 3 |  |
|  |  | 3 2 |  |

For both undirected and directed cases, in both time and frequency domains, we compute three types of measure:

**SRC**: Per-source *unconditional* measures, which measure associations between a single source and all remaining sources,

**ROI**: Per-ROI *pairwise-conditional* measures, which measure associations between pairs of ROIs, conditioned on all other sources, and

**GLO**: Intra-ROI *global-conditional* measures, which measure the total statistical association of all sources within an ROI, conditioned on all sources lying outside the ROI.

The GLO measures may be interpreted in terms of “density” of within-ROI statistical associations, in the spirit of (undirected) multi-information (Studený and Vejnarová, 1998) and (directed) causal density (Seth, 2009; Seth et al., 2011), or global transfer entropy (Barnett et al., 2013).

#### 2.2.1 Undirected measures: partial correlation and partial coherence

Given jointly-distributed multivariate random variables *X, Y, Z*, we have the conditional mutual information *I* (*X* : *Y*|*Z*). Mutual information (Cover and Thomas, 1991) quantifies the degree to which two jointly-distributed variables are statistically dependent. In the case that *X* and *Y* are univariate and *X, Y, Z* are jointly multivariate-normally distributed, *I* (*X* : *Y*|*Z*) = log [1 − *ρ*(*X, Y|Z*)^2^], where *ρ*(*X, Y*|*Z*) is the conventional partial correlation coefficient, so that *I* (*X* : *Y*|*Z*) is a monotonic function of the (squared) partial correlation^3^; in this sense *I* (*X* : *Y*|*Z*) generalises partial correlation to multivariate (and non-Gaussian) variables. Under Gaussian assumptions:

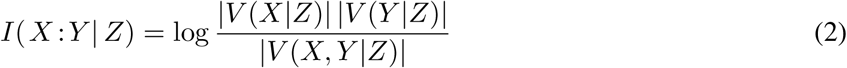

where *V* (*X*|*Z*), *V* (*X, Y*|*Z*), etc., denote partial variance-covariance matrices, |···| denotes a determinant, and “log” always denotes natural logarithm.

If *X* = {*X*_*i*_} is multivariate, a conditional version of multi-information can be defined as *I* (*X*|*Z*). Multi-information (Studený and Vejnarová, 1998) quantifies the degree to which a jointly-distributed set of variables are mutually dependent. Under Gaussian assumptions, multi-information corresponds to the “partial total correlation”:

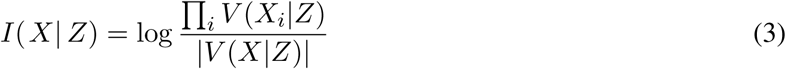

which is the basis for the (undirected) global-conditional measure we use (GLO).

In the frequency domain, we use 2nd-order statistics as for the time-domain Gaussian case above, but based on the cross-power spectral density (CPSD) matrices *S*(*X*; *ω*), etc., rather than the covariance matrices *V* (*X*), where *ω* denotes frequency (in Hz) on the range *ω* ∈ [0, *ν/*2] with *ν* the sampling frequency. We thus consider the measure

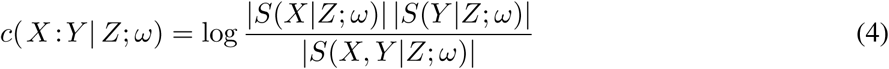

In case *X, Y* are univariate, *c* (*X* : *Y*|*Z*; *ω*) is the logarithm of the standard partial coherence measure. For multivariate processes, we also have the global-conditional spectral measure [*cf*. (3)]

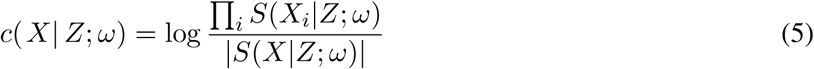

#### 2.2.2 Directed measures: Granger causality in time and frequency domain

Directed connectivity measured are based on Granger causality (GC) (Wiener, 1956; Granger, 1963, 1969; Geweke, 1982), a 2nd-order statistic for stochastic time-series based on optimal linear prediction. We use Geweke’s multivariate conditional form (Geweke, 1984) in time and frequency domains. Although the mathematics of GC have been well described previously [see, e.g., Barnett and Seth (2014)], we specify our derivation below in order to prevent ambiguity in interpretation, especially for the more complex cases of conditional GC in the frequency domain.

Let *X*_*t*_ (target), *Y*_*t*_ (source) and *Z*_*t*_ (conditioning variable) be jointly-distributed, wide-sense stationary (possibly multivariate) processes. The optimal linear predictor (in the least-squares or maximum-likelihood sense) of *X* based on the past histories of *X, Y, Z* is given by the conditional expectation 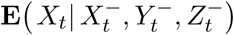, where 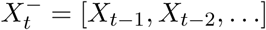, etc., denotes the history of a process up to the previous time step. The residual prediction errors are then 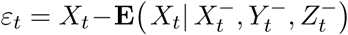; we denote the covariance matrix of residual errors— a measure of predictive efficacy—by 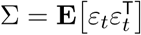. Now consider the optimal predictor 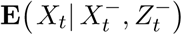 of *X* based on the histories of *X* and *Z* alone; i.e., omitting the source variable *Y*. The residuals of this “reduced” predictor can be written as 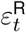, and its covariance matrix as 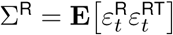. Then the (time-domain) GC from source *Y* to target *X* conditional on *Z* is defined to be

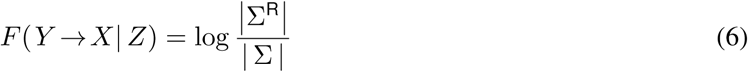

and may be interpreted intuitively as “the degree to which *Y* predicts the future of *X*, over and above the degree to which *X* already predicts its own future, controlling for *Z*”. In finite sample, the infinite histories 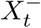, etc., are replaced by [*X*_*t−*1_, …, *X*_*t−p*_], etc, where *p* is the selected model order [*cf*. (1)], and the statistic (6) becomes a log-likelihood ratio.

Since GC, as a directed measure, is not symmetrical in the source and target variables, we have an inbound and an outbound version of the unconditional form (SRC), corresponding respectively to system → source and source → system information flow. For multivariate processes *X*_*t*_ = {*X*_*ti*_}, analogous to conditional multi-information in the undirected case, a global-conditional measure (GLO) may be defined as the sum of conditional causalities from *X* itself to each individual variable *X*_*i*_ (Barnett et al., 2013):

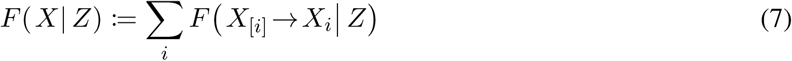

where subscript [*i*] denotes omission of the *i*th index.

Geweke also introduced a spectral (frequency domain, conditional) GC statistic, written here as *f*(*Y*→*X*|*Z*;*ω*); for details see Geweke(1982, 1984). A corresponding global-conditional measure *f*(*X*|*Z*; *ω*) can be defined as ∑_*i*_ *f*(*X*_[*i*]_ →*X*_*i*_ |*Z*; *ω*).

For empirical data, we calculate GC in time and frequency domain from the estimated VAR parameters using a state-space method (Hannan and Deistler, 2012; Barnett and Seth, 2015; Solo, 2016)^4^.

#### 2.2.3 Frequency-band averaging

The relationship between time- and frequency-domain measures is underpinned by Szegö’s Theorem, which yields (Rozanov, 1967):

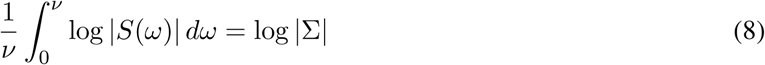

where *ν* is the sampling frequency. As a corollary, spectral GC integrates across the broadband spectrum to yield the corresponding time-domain measure (Geweke, 1984):

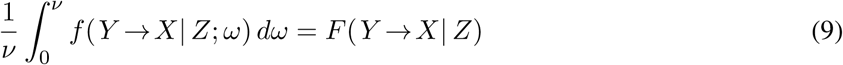

This relation thus holds for the directed spectral measures^5^. In general, all the frequency-domain measures (undirected and directed) can be integrated (averaged) over a frequency band [*ω*_1_, *ω*_2_] to yield a “band-limited” measure (Barnett and Seth, 2011) 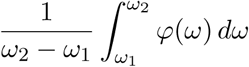, where *φ*(*ω*) represents any of our spectral measures. All frequency-domain results in Section 3 are presented as band-limited over the frequency bands in Table 3.

### 2.3 Statistical inference

All time-domain 2nd-order statistics take the form of log-likelihood ratios in finite sample, so that the classical large sample theory (Neyman and Pearson, 1928; Wilks, 1938; Wald, 1943) applies, yielding asymptotic *F* or *χ*^2^ distributions for their sample estimators. Sampling distributions for the frequency-domain statistics are generally not known analytically; we derive them empirically by subsampling/surrogate data methods. We used a False Discovery Rate (FDR) correction to correct for multiple hypotheses per-drug/subject/condition/measure (Benjamini and Hochberg, 1995). For the directed FC measures, variation of empirical model order per-drug/subject may introduce a source of bias; however, given the large number of sample observations per subject/condition (i.e., number of epochs × MEG observations per epoch), bias and variance, estimated from the corresponding *χ*^2^ distribution under the null hypothesis, were an order of magnitude smaller than typical between-condition changes in the statistical measures, and could thus be safely ignored. Cross-subject statistical comparisons of FC measures between conditions (drug vs placebo) were carried out using Wilcoxon’s signed rank-test (Wilcoxon, 1945) (paired t-tests could not be used since sample estimators are non-Gaussian). The rank correlation, defined as Wilcoxon’s W-statistic normalised by the total rank sum to lie between −1 and +1, is presented as an effect size in all results. Significance is presented at *α* = 0.05, with FDR multiple-hypothesis adjustment.

## 3 Results

### 3.1 Spectral power

Before analysing FC, we first examine cross-subject changes in spectral power between drug and placebo. For LSD, psilocybin (PSI), and ketamine (KET), results (Figure 1) are broadly consistent with Muthukumaraswamy et al. (2013) [PSI] and Carhart-Harris et al. (2016) [LSD], and reveal a similar effect for sub-anesthetic ketamine. For LSD, PSI, and KET there is a general decrease in spectral power across all frequency bands, but especially in the *δ*–*β* bands. For tiagabine (TGB), by contrast, there is a slight increase in spectral power in *δ*−*α*. Figure 2 plots spectral power for the 90 sources for a representative single subject for each drug. For the psychedelics, there is a decrease in spectral power in the *δ*–*β* range against an approximately 1*/f* background, and, in line with previous analyses (Muthukumaraswamy et al., 2013), there is also a slight shift of the *α* peak to a higher frequency, most obviously for LSD [see Muthukumaraswamy and Liley (2018)].

**Figure 1:**
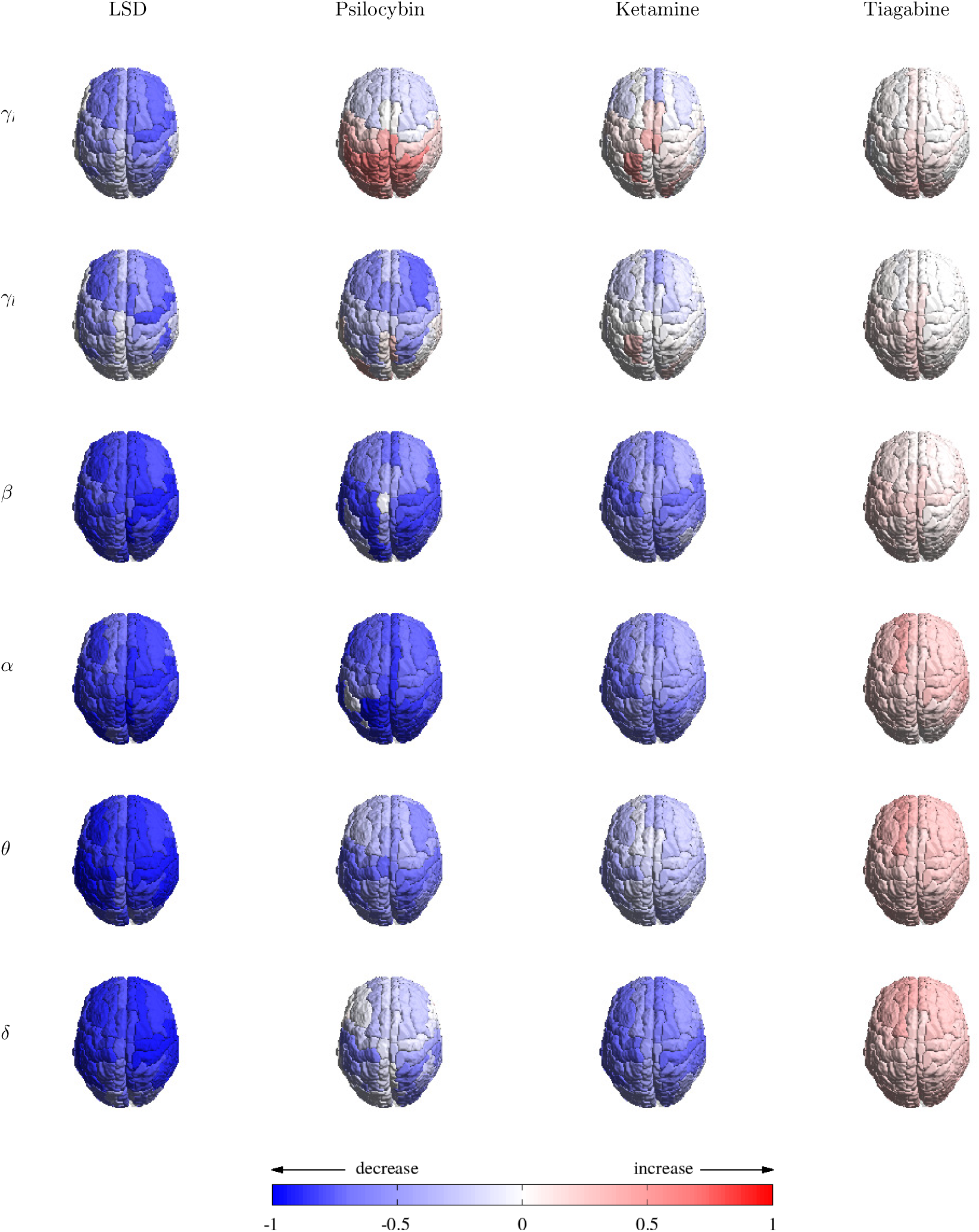
Change in spectral power by source and frequency band. Spectral estimates are based on a multi-taper method averaged over epochs (Section 3.2). In this and subsequent figures, colour indicates cross-subject effect size (rank correlation; see Section 2.3): red indicates an increase in the measured quantity for drug vs placebo, blue a decrease.

**Figure 2:**
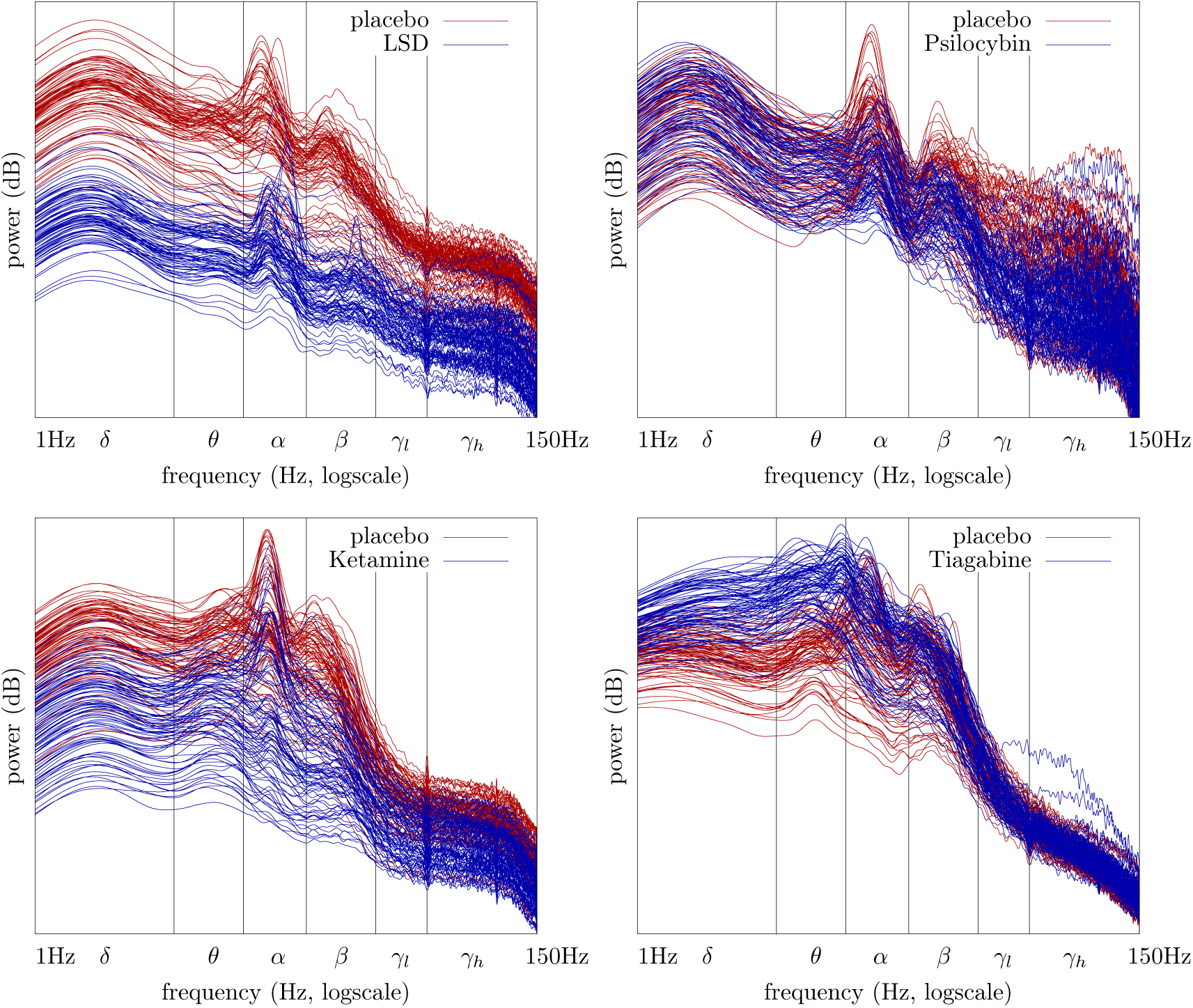
Spectral power for representative single subjects: drug (blue) vs. placebo (red). Lines plot auto-power for the 90 sources.

### 3.2 Functional connectivity

We next examine changes in per-source undirected (MI) and directed (GC) time-domain FC for drug vs placebo. Figure 3 shows changes in the unconditional per-source measures (SRC): undirected (top row), directed inbound (middle row) and directed outbound (bottom row).

**Figure 3:**
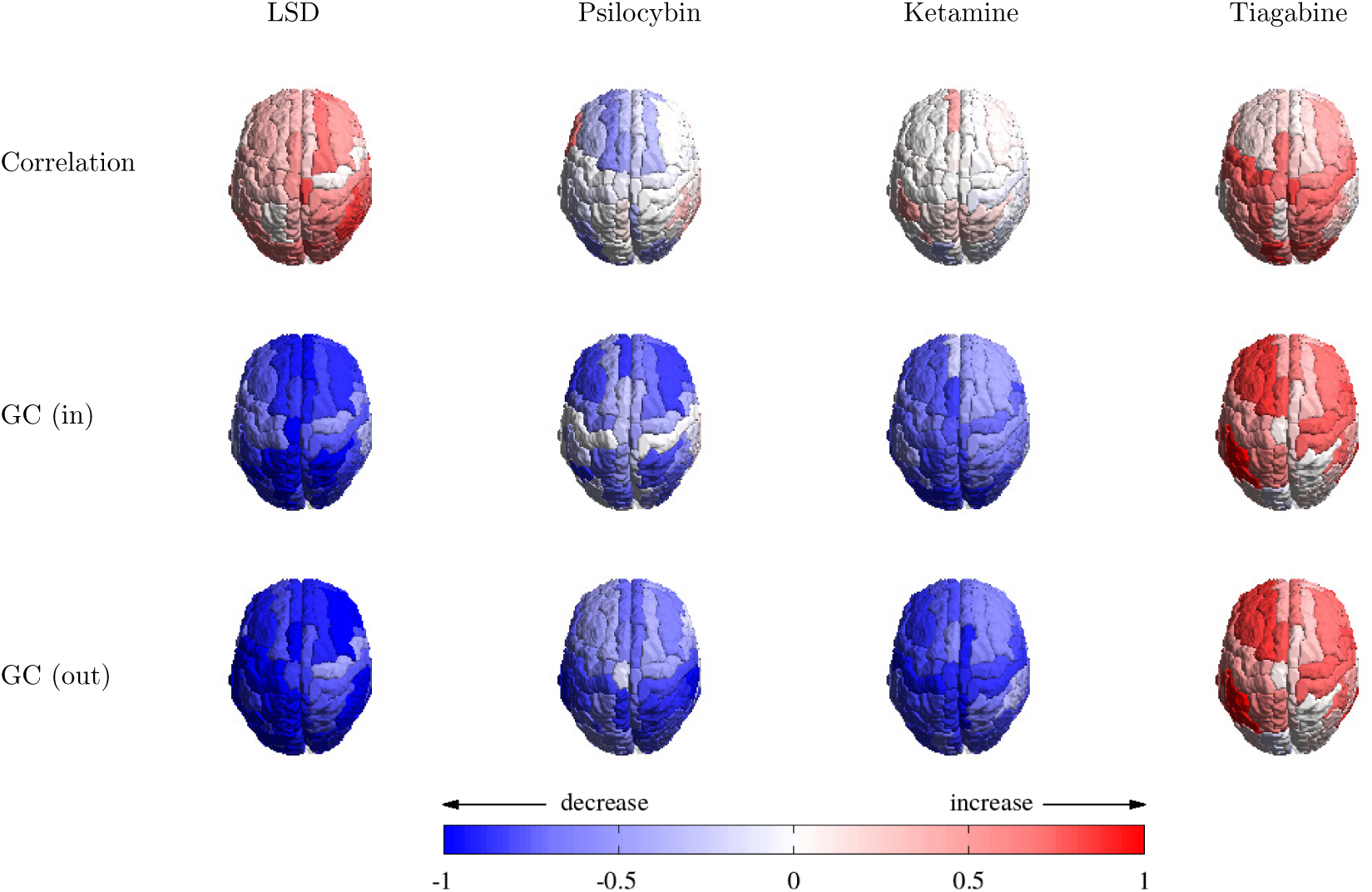
Time domain, unconditional (SRC): change in undirected (MI) vs. change in directed (GC) FC measures between single source and rest of brain. Top row: undirected; middle row: directed, inbound; bottom row: directed, outbound.

For the directed measures, inbound and outbound, there are significant decreases in FC across all the psychedelics (as compared with placebo), while for tiagabine there is an increase. Results for undirected measures are less clear: for LSD and tiagabine there are significant increases in MI, while for psilocybin and ketamine results are inconclusive. Notably, within the psychedelics, effect sizes for LSD are generally stronger than for psilocybin or ketamine.

Figures 4-6 decompose the per-source time-domain results by frequency. For the undirected case, Figure 4 shows that the increase in MI in LSD is strongest in the *γ*-band (but see Section 4.4). While results for psilocybin and ketamine are statistically weak, there is a suggestion of a slight increase in *γ*. For tiagabine the increase is strongest in *δ* and *θ*. These observations suggest a distinct mechanism of action underlying increases in MI in tiagabine as compared with the psychedelics. (We interpret the *γ*-band results cautiously, given potential for confounds due to muscle artefact; see Section 4.4.)

**Figure 4:**
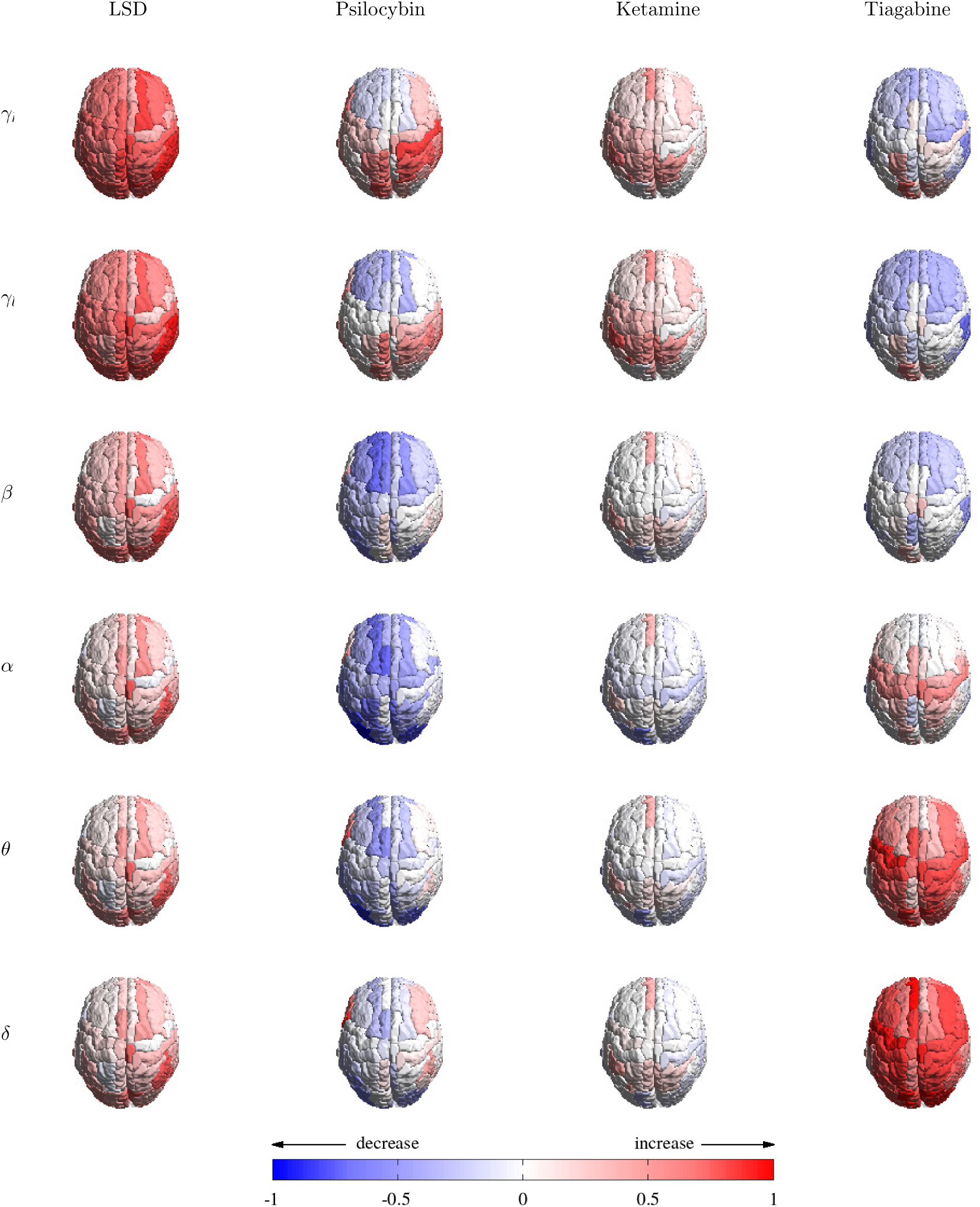
Frequency domain, unconditional (SRC): change in undirected FC (MI) between single source and rest of brain.

**Figure 5:**
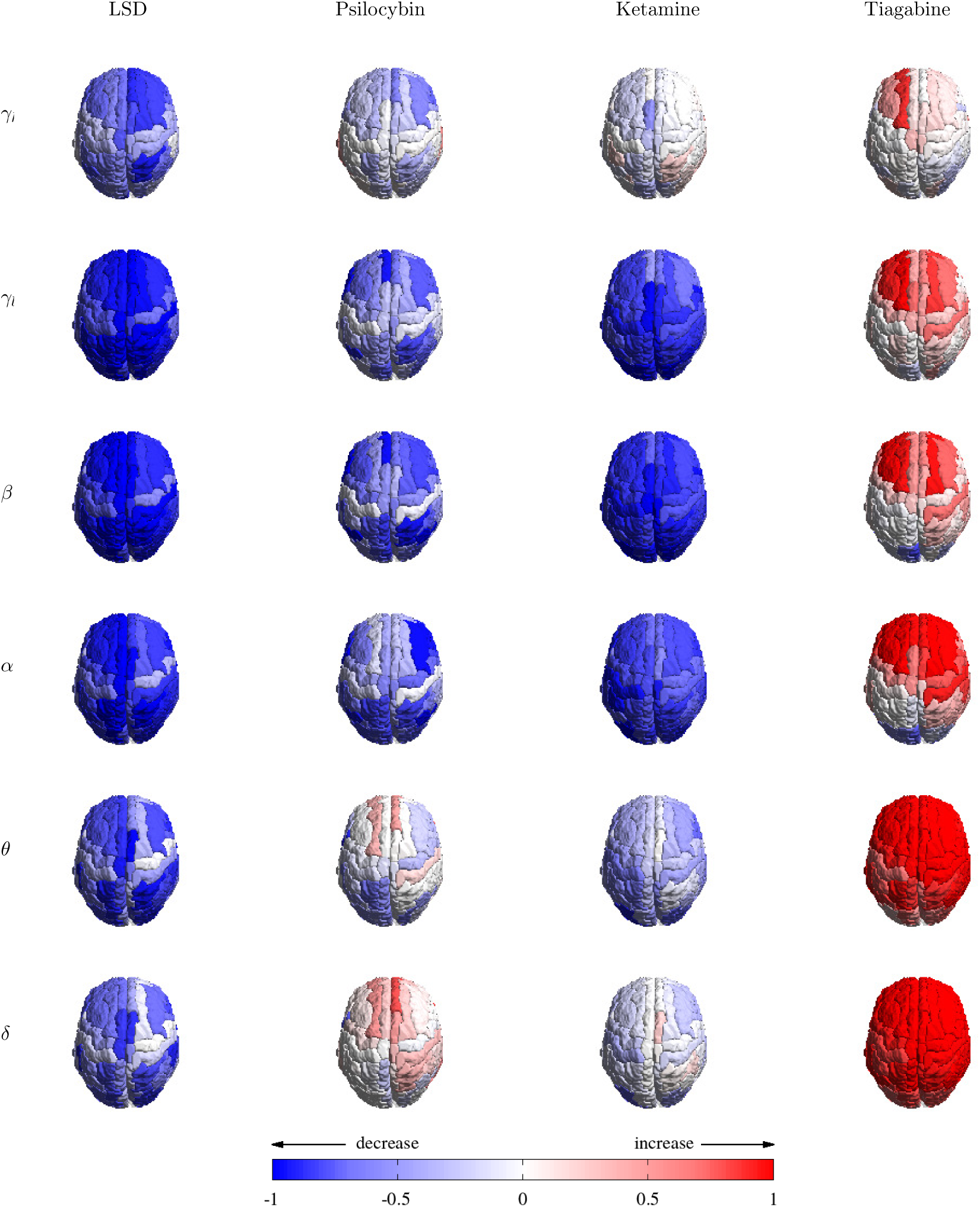
Frequency domain, unconditional (SRC): change in directed (inbound GC) FC between single source and rest of brain.

**Figure 6:**
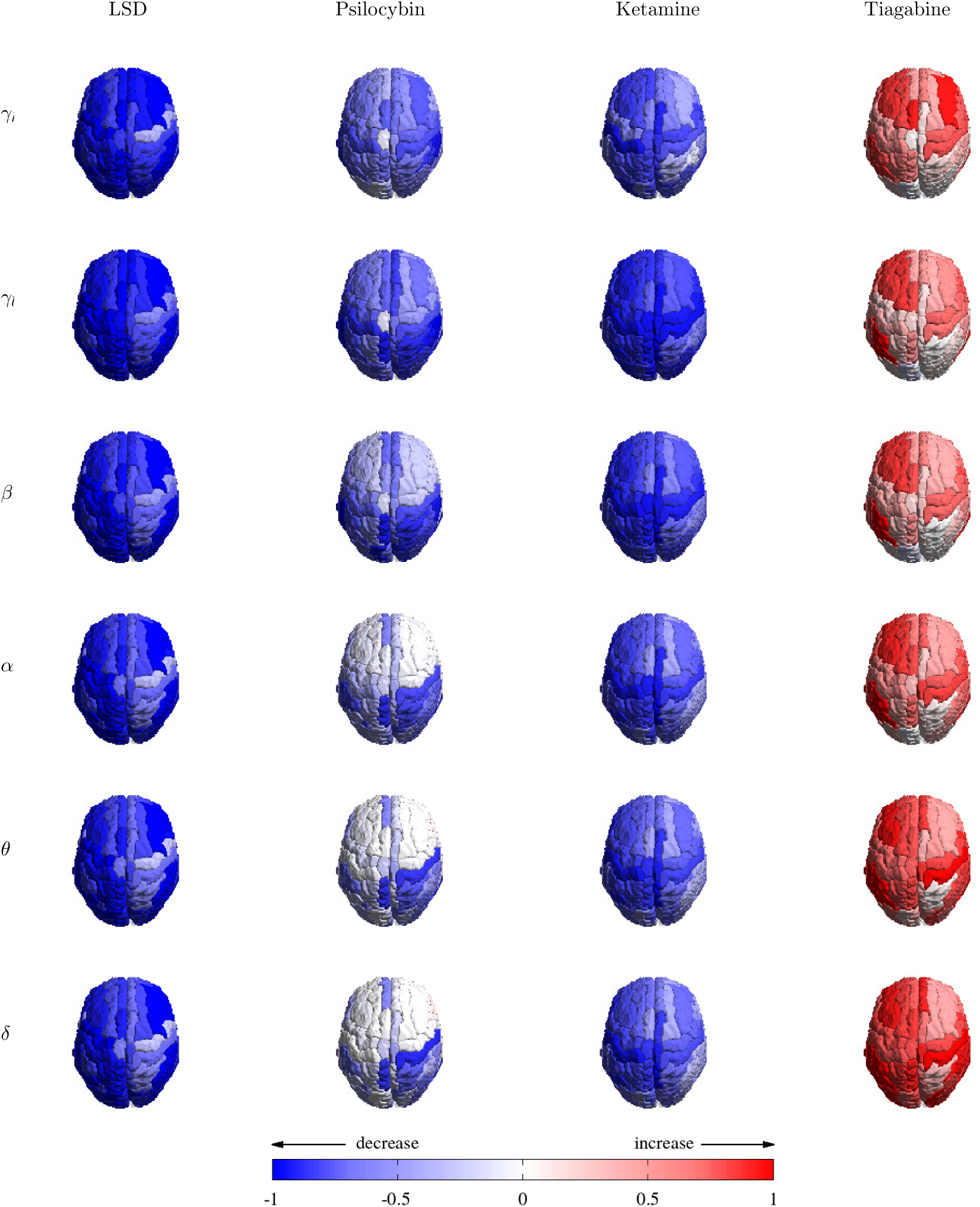
Frequency domain, unconditional (SRC): change in directed (outbound GC) FC between single source and rest of brain.

Figures 5 and 6 reinforce the time-domain results for changes in the directed measures. For the psychedelics, the decrease in GC is broadband, particularly for LSD (for psilocybin the decrease is stronger in *α* − *γ*_*l*_). For tiagabine, the increase in inbound GC is strongest in *δ* − *θ*, while for outbound GC it is evenly spread across the spectrum.

We next examined the regional specificity of changes in directed and undirected FC. Figure 7 presents results for the inter- and intra-ROI measures (ROI and GLO). The left-hand column displays changes between drug and placebo for undirected (MI) measures. The right-hand column displays results for the directed (GC) measures. For MI, differences for psilocybin and ketamine are again small (*cf*. Figures 3 and 4). For LSD, the increase in MI is strongest between the occipital region and other ROIs, particularly the cingulate region. For tiagabine, the increase in MI is pronounced between the parietal and frontal/limbic/occipital regions, again suggesting a distinct mechanism. Intra-ROI changes in MI are weaker than inter-ROI changes. Turning to directed measures, for LSD decreases in GC are cortex-wide, both within and between ROIs. For psilocybin, the decrease is strongest between the parietal and other regions, although intra-region increases are only slight. For ketamine, the strongest decreases are from other ROIs to the parietal and occipital regions, with significant intra-ROI decreases in occipital, parietal and sensorimotor regions. For tiagabine, the increase in GC is fairly evenly spread across the cortex, with the exception of the occipital region (there is also a significant decrease in intra-ROI directed connectivity in the occipital region).

**Figure 7:**
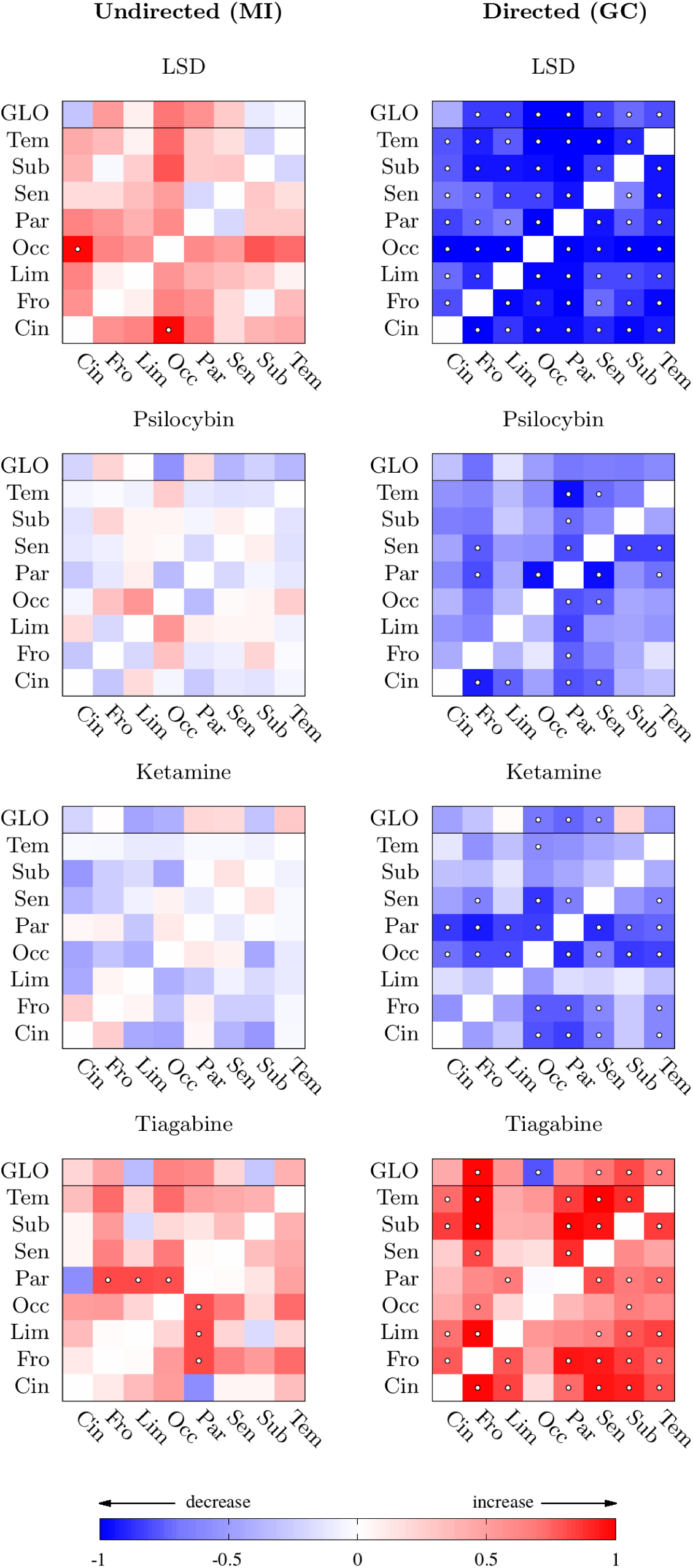
Time domain, inter-ROI pairwise-conditional (ROI) and intra-ROI global-conditional (GLO): change in undirected vs. change in directed measures between ROIs (Table 1), conditioned on rest of brain. The top row of each panel (GLO) shows effect sizes per column for the global-conditional intra-ROI measures (GLO). The undirected plots are symmetrical; for the directed plots, row label references “to” ROI, column label “from” ROI. White dots indicate a statistically significant cross-subject effect at *α* = 0.05, Wilcoxon signed-rank test, with FDR correction (see Section 2.3).

Summarising, comparing psychedelics with placebo reveals significant decreases in GC, widespread across cortical regions and across frequency bands, and which are most pronounced for LSD. These decreases in directed FC contrast with either little change, or increases, in undirected FC as measured by MI. The non-psychedelic GABA reuptake inhibitor tiagabine shows increases in both GC and MI, compared with placebo.

To verify that these results reflect meaningful changes in directed and undirected FC, we conducted further analyses to explore whether the observed changes in MI and GC could be accounted for by (i) changes in power spectra, including effects of signal-to-noise ratio (SNR), and (ii) changes in residuals correlation in the VAR modelling of the data.

### 3.3 Relation between changes in undirected and directed FC

A novel feature of our analysis is that we compare GC and MI on the same data, and we see in our empirical results a variety of ways in which they relate to each other. Thus (Figure 3) we see the striking opposite movement for LSD, same-direction movement for TGB, while for the other psychedelics (PSI and KET) the decrease in GC is not obviously accompanied by increases in MI. To better understand the extent to which GC and MI are reflecting distinct aspects of neural dynamics, we conducted further theoretical, modelling, and empirical analyses.

#### 3.3.1 Spectral power and signal-to-noise ratio

Existing functional analyses of the psychedelic state (Riba et al., 2004; Muthukumaraswamy et al., 2013; Carhart-Harris et al., 2016; Pallavicini et al., 2019) have concentrated on changes in spectral power (Section 3.1). We therefore wondered whether the decreased directed functional connectivity we observed might have a straightforward explanation in terms of changes in spectral power. This possibility can be reasonably rejected for the following reason: previously conducted power analyses have focused on auto-spectral power, and have revealed generally broadband decreases in the psychedelic state - as we also find (Figures 1, 2). Such broadband changes most likely reflect a rescaling of neural signals (but see below). Critically, Granger causality is scale invariant (Barrett and Barnett, 2013). In fact, more generally, Granger causality is invariant under (invertible) filtering (Geweke, 1982; Barnett and Seth, 2011). Therefore, changes to auto-spectra are not by themselves informative about how GC might be expected to change. Because GC depends on the full cross-power spectrum—in complicated ways (Dhamala et al., 2008)—our results cannot be readily accounted for in terms the observed changes in broadband auto-spectra.

Another possibility is that the decrease in directed FC between placebo and drug conditions might be due to changes in signal-to-noise ratio (SNR). Previous studies (Muthukumaraswamy et al., 2013; Carhart-Harris et al., 2016) indicate that the psychedelic drugs are associated with a decrease in SNR; this is corroborated by our results for spectral power changes in the psychedelic conditions (Figures 1, 2). To test whether a reduction in SNR might account for a reduction in directed FC, we investigated whether the addition of correlated, additive broadband white noise [representing a mixture of highly correlated room noise plus uncorrelated sensor noise (Vrba and Robinson, 2001)] to the placebo data could reproduce the time-domain FC results. Results, illustrated in Figure 8, demonstrate that a sufficient level of additive noise (≈20 dB) to roughly emulate the decrease in GC seen in Figure 3, also results in a strong *decrease* in MI^6^, in direct contrast to Figure 3. We thus conclude that a decrease in SNR in the psychedelic state is highly unlikely to account for the observed changes in FC.

**Figure 8:**
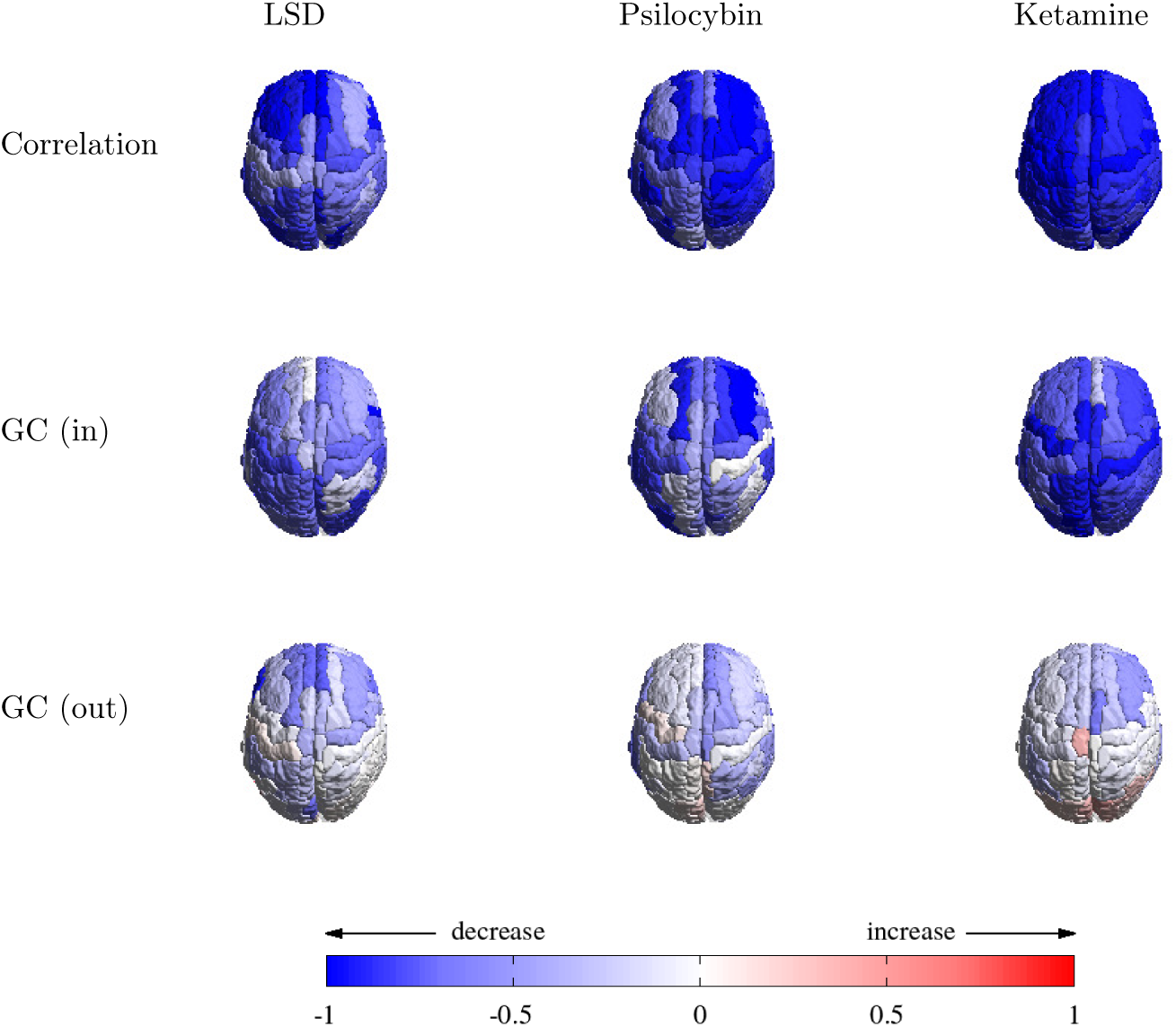
Time domain, unconditional (SRC): change in undirected vs. change in directed FC measures between single source and rest of brain (psychedelic drugs) for placebo vs. placebo+additive, correlated noise. Top row: undirected; middle row: directed, inbound; bottom row: directed, outbound. Additive noise was at a level of ≈ 20 dB, with a Pearson correlation coefficient of *ρ* ≈ 0.9 between sources.

#### 3.3.2 Empirical and theoretical relationship between GC and MI

The opposite movement of GC and MI seen in LSD—as well as the same-direction movement in tiagabine— are striking and deserve further analysis. One might think, *a priori*, that directed and undirected measures of FC should move together: that increased MI might imply increased information flow (as measured by GC). However, formally, mutual information and information flow are not directly related. That is, there is no theoretical reason to expect that a decrease (due to LSD, for example) in GC between two ROIs should be associated with an increase in MI, or vice-versa.

To probe the relationship between GC and MI in our data, we firstly performed the following empirical analysis. For each drug we investigated the correlation across subjects, between ΔGC = GC(drug) − GC(placebo) and ΔMI = MI(drug) − MI(placebo), both inter- and intra-ROI. That is, we asked empirically whether larger changes in undirected measures corresponded with larger changes in directed measures. As Figure 9 shows, there is no evidence of consistent correlation between ΔGC and ΔMI. Indeed, for specific ROIs, change in the measures were sometimes found to be correlated and sometimes anti-correlated – but statistical significance was in any case generally negligible.

**Figure 9:**
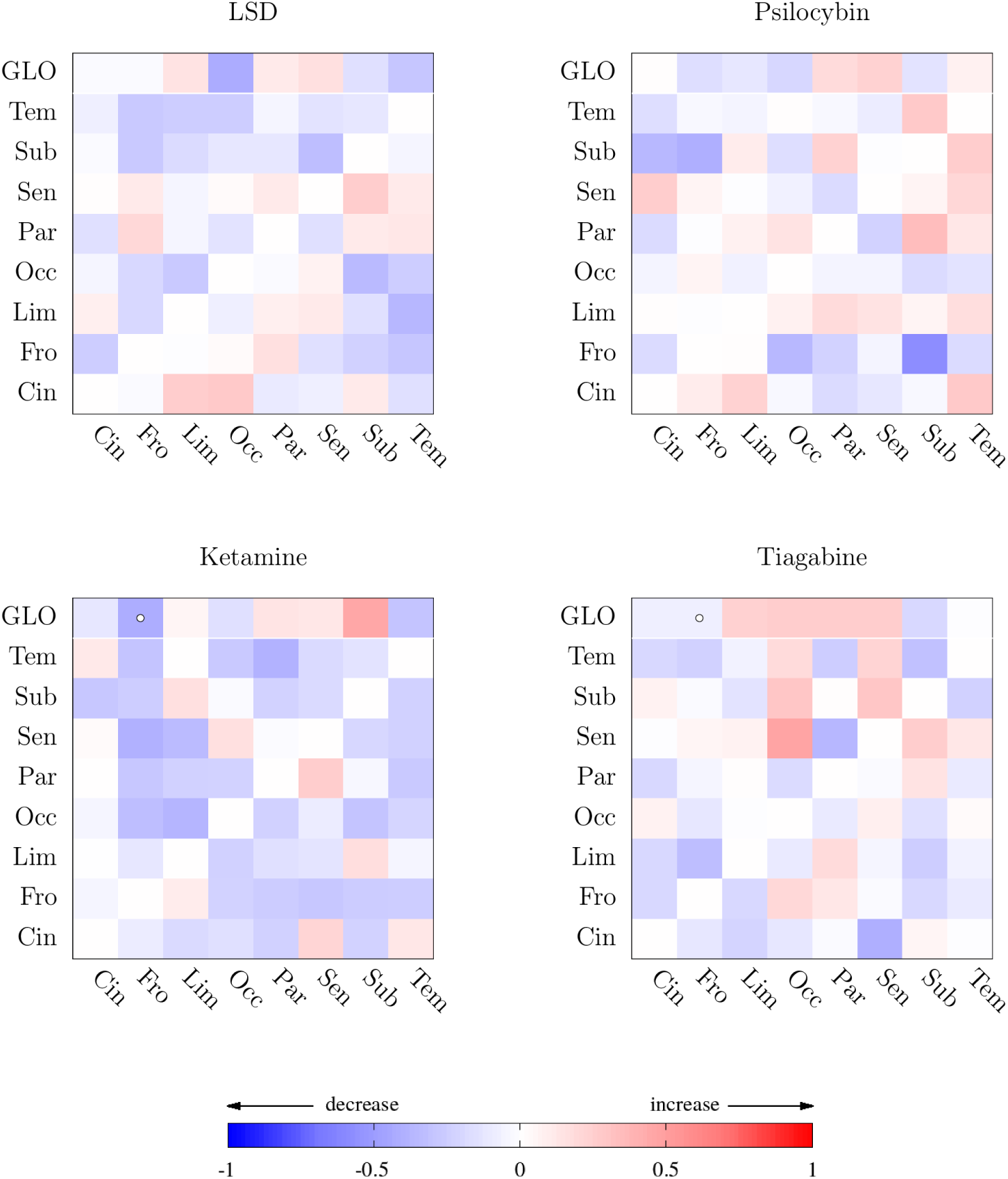
Inter- and intra-ROI correlation between ΔGC = − GC(drug) − GC(placebo) and ΔMI = MI(drug) MI(placebo). Correlation is calculated, per ROI pair (intra, ROI) or ROI (intra, GLO) as Kendall’s *τ* rank-correlation statistic, in range [−1, 1]. Small circles indicate statistical significance at 95% confidence, with a per-plot false discovery rate (FDR) correction for multiple hypotheses.

To further examine this issue we conducted a theoretical analysis, examining the relationship between MI and GC for the stationary bivariate, VAR(1) model

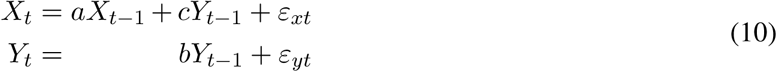

with unidirectional GC and residuals correlation *ρ*(*ε*_*xt*_, *ε*_*yt*_) = *κ*; see A for details.

Figure 10a and Figure 10b display heat maps of MI *I*(*X* : *Y*) and GC *F* (*Y* →*X*) for *κ* = 0, fixed *a*, and *b, c* varying within the model parameter space. The green arrows indicate the direction of steepest gradient ∇*I*(*X* : *Y*), while the magenta arrows indicate the direction of steepest gradient ∇*F* (*Y* →*X*) for GC. At a given point in parameter space, the angle *θ* between the gradients is given by (A)

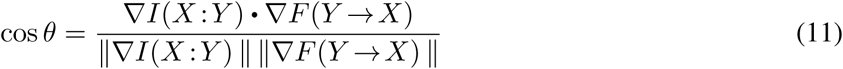

The cosine is +1 when the gradients are in exactly the same direction, 1 when they point in opposite directions and 0 when they are orthogonal (Figure 10c). For *κ* = 0, the gradients never point in opposite directions, but (Figure 10d-f) this is not necessarily the case for *κ* ≠ 0. Together, these figures show that there will always be directions in parameter space where MI and GC will be changing either in the same, or in opposite, directions. Analysis of this minimal VAR model therefore further establishes that GC and MI can move independently under variations in the underlying data generating process.

**Figure 10:**
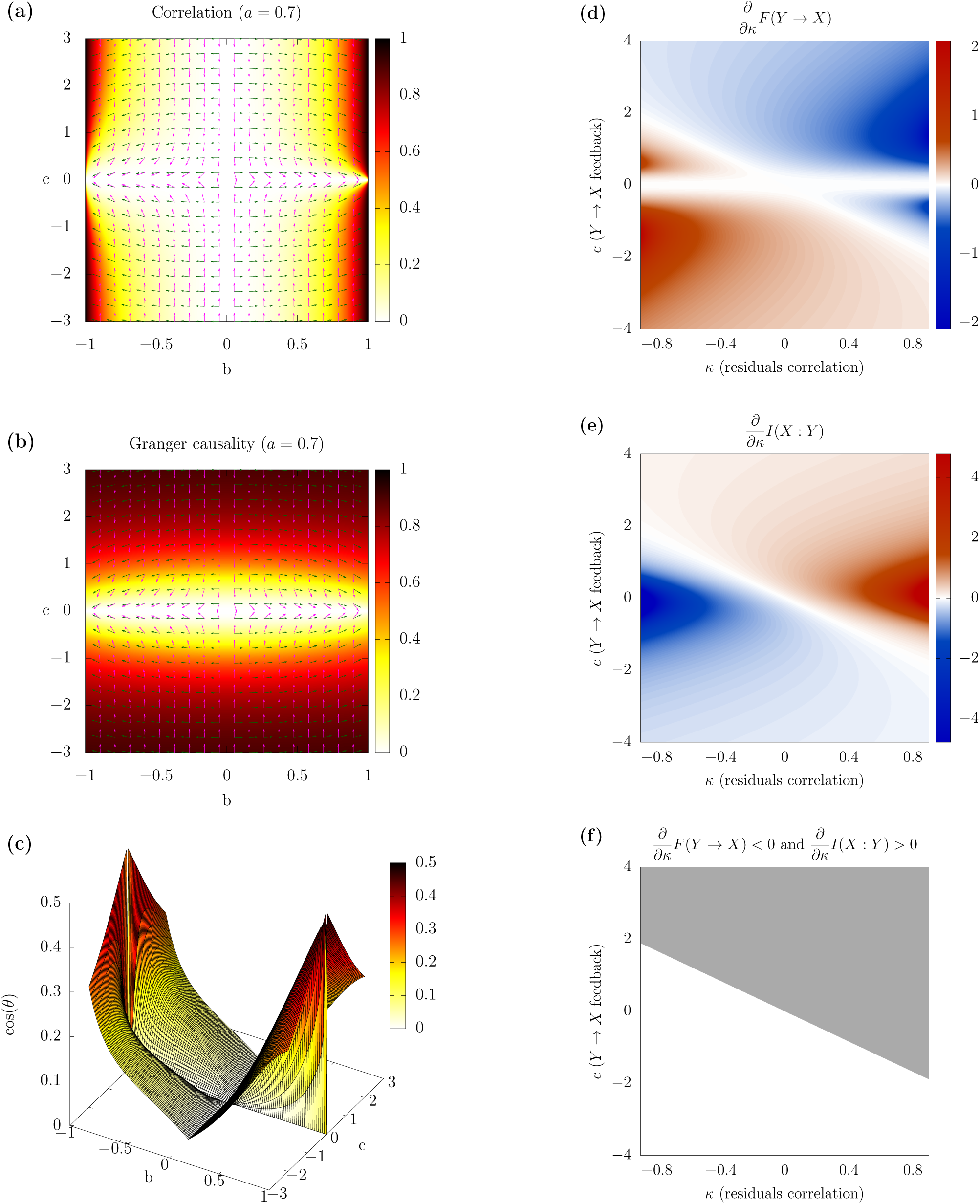
Left column: heat maps of *I*(*X* : *Y*) **(a)** and *F* (*Y* → *X*) **(b)** for the VAR model (10) with *κ* = 0, for fixed *a* and *b, c* varying over parameter space. Green arrows indicate the direction of steepest gradient ∇*I*(*X* : *Y*), while the magenta arrows indicate the direction of steepest gradient ∇*F*(*Y* → *X*). Fig. **(c)** displays a surface plot of cos *θ*, where *θ* is the angle between the gradients of *I*(*X* : *Y*) and *F* (*Y* → *X*) with *κ* = 0, for fixed *a* and *b, c* varying over parameter space. cos *θ* is +1 when the gradients point in the same direction, −1 when they point in opposite directions, and 0 when they are orthogonal. Right column: heat maps of the gradient of *F* (*Y* → *X*) **(d)** and *I*(*X* : *Y*) **(e)** with respect to *κ* plotted against causal strength *c* and residuals correlation *κ*, for *a* = 0.3, *b* = 0.4. In Fig. **(f)**, the region where *I*(*X* : *Y*) increases and *F* (*Y* → *X*) decreases with increasing *κ* is marked in grey.

Given that GC and MI (i) *need not* change together, as established theoretically and via modelling, and (ii) generally *do not* move together—at least not consistently—as shown by the heterogeneity in the relation between GC and MI across the different drugs (Figure 9), what could explain the striking opposite movement in the specific case of LSD?

One possibility relates to correlations between the residuals in the corresponding VAR models^7^. In our minimal VAR model, the behaviour of the MI and GC gradients as residuals correlation *κ* changes is striking. In A we derive explicit expressions for 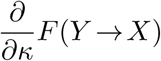 and 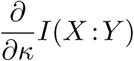. Extensive simulations show that, for any values (except *c* = 0) in the (*a, b, c, κ*) parameter space, the signs of these quantities are always opposite. Thus any change in residuals correlation always results in an opposite movement of MI and GC. For at least half of the entire parameter space (in the minimal VAR model), increasing *κ* (i.e., increasing residuals correlation) always leads to increasing MI coupled with decreasing GC, while for the other half, increasing *κ* leads to decreasing MI coupled with increasing GC (Figure 10, right column). Therefore, even though a change in residuals correlation is consistent with the empirical changes in MI and GC observed with LSD, it is less consistent with PSI (where an increase in residuals correlation accompanies a decrease in GC but no significant change in MI), and not at all with KET (where there is no change in residuals correlation), or with TGB (where a decrease in residuals correlation leads to increases in both GC and MI). These analyses therefore indicate that changes in residuals correlations cannot readily explain the pattern of empirical results across all drugs.

Figure 11 displays drug vs. placebo change in residuals correlation for the VAR models (i.e., the models fitted to the MEG data, not the minimal VAR model discussed above). For LSD and psilocybin, there is a statistically significant increase in residuals correlation in the drug condition, while for tiagabine there is a statistically significant decrease. On the assumption that the minimal VAR model result—that *any* change in residuals correlation between conditions leads to the opposite movement of directed and undirected FC— generalises to more complex network scenarios, it is unlikely that changes in residual correlation can explain the relationship between directed and undirected FC that we observe across all drugs. (In addition, while for KET there are hints of an increase in undirected FC to accompany the decrease in GC, there are no significant changes in residuals correlation). Therefore, alterations in residuals correlation might account for the pattern of results for LSD but cannot account for the pattern in general. We have not established the extent to which our minimal VAR result generalises; but even in that model there are other possible structural mechanisms which may drive the effect which do not involve residuals correlation (*cf*. Figure 10, left column).

**Figure 11:**
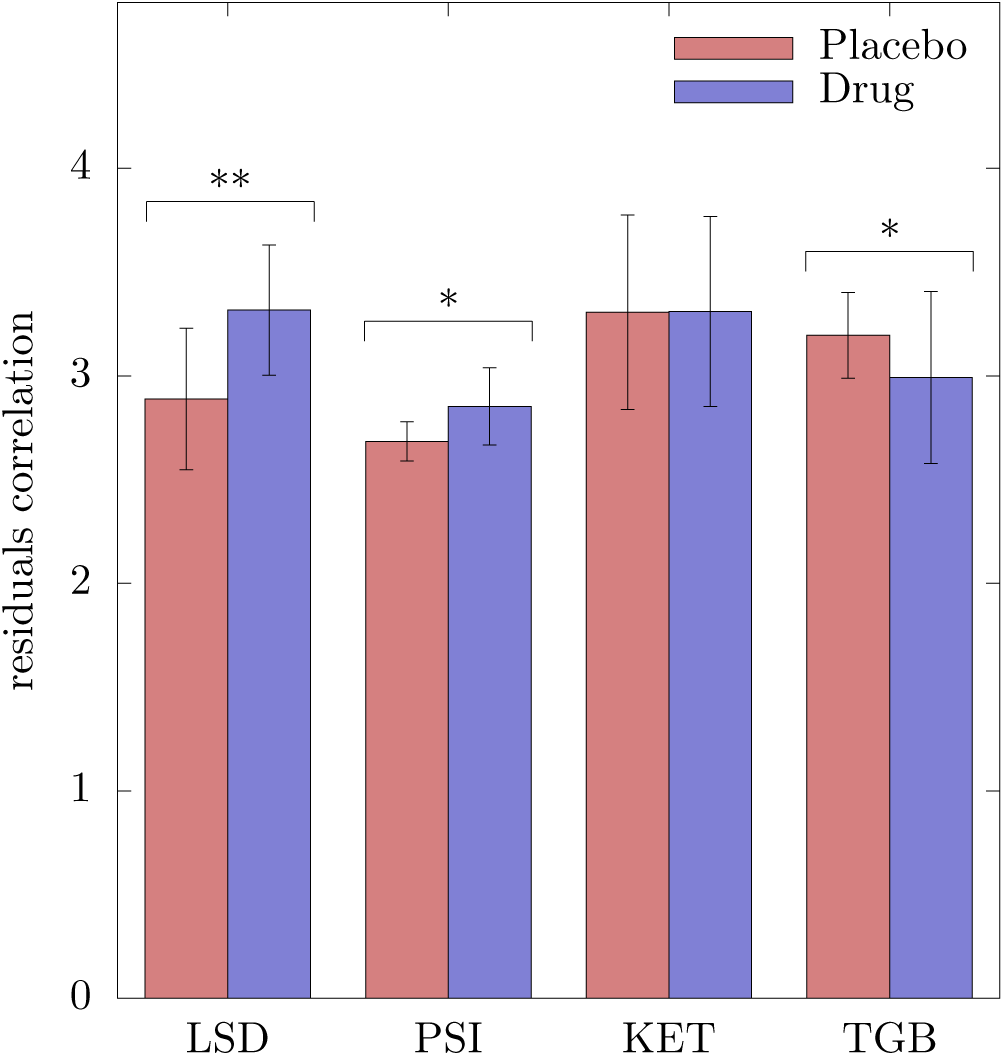
VAR residuals correlations, measured as multi-information of the residuals normalised by the multi-information of a uniform random correlation matrix of the same size. The figure plots median residual correlations across subjects, with error bars at *±*1 mean absolute deviations.

Altogether, the mechanistic processes underlying the relative changes in directed and undirected FC in the different drug conditions remain to be fully elucidated. Importantly, though, the changes we observe cannot readily be explained away in terms of changes in power auto-spectra, signal-to-noise ratio, or residuals correlation.

## 4 Discussion

In this exploratory study, we examined the effects of three psychedelic drugs on large-scale brain dynamics in terms of directed functional connectivity (FC). Unlike more familiar undirected FC measures, such as correlation and coherence, directed FC measures take into account temporal dependencies in the data, potentially delivering fresh insights into alterations to neural dynamics, and more specifically into changes in information flow both between and within brain regions.

We applied both directed FC measures (Granger causality/information flow) and undirected measures (correlation/mutual information), in time and frequency domains, to source-localised MEG data obtained in resting state conditions, contrasting placebo against three different psychedelics (LSD, psilocybin, low-dose ketamine), as well as against a non-psychedelic control, tiagabine. Our main result revealed a consistent broadband decrease in information flow in psychedelic conditions, both between and within brain regions, broadly across cortex. In the case of LSD, this decrease in information flow was accompanied by an increase in undirected FC. By contrast, for the tiagabine control, we observed increases in both directed and undirected FC in the drug condition.

Further empirical and theoretical analyses examined whether these changes in FC could be accounted for by changes in power spectra, signal-to-noise ratio, or correlation between the residuals of the predictive VAR models used to derive FC statistics. We verified that the observed changes in directed and undirected FC could not readily be accounted for by these factors.

### 4.1 Power spectra

We first identified a substantial reduction in broadband spectral power in the psychedelic state. This is a relatively well established effect, which has been described in several previous studies: see, e.g., Fink (1969) (LSD, mescaline/EEG), Riba et al. (2004) (ayahuasca/EEG), Muthukumaraswamy et al. (2013) (psilocybin/MEG) and Carhart-Harris et al. (2016) (LSD/MEG). Recently, Pallavicini et al. (2019) analysed spectral changes in LSD, psilocybin and ketamine using the same MEG dataset as used in this study. They report region-specific patterns of spectral power reduction in the alpha and theta bands common to LSD and psilocybin, as well as changes in the beta band common to all three drugs. Our spectral analyses (Figure 1) are consistent with these findings, as would be expected given we analyse the same data. Interestingly, accompanying these broadband reductions in power, we also note a (small but significant) upward shift in the alpha peak frequency (Figure 2), which is most evident for LSD (Walter, 1957; Carhart-Harris et al., 2016; Muthukumaraswamy and Liley, 2018). The mechanisms and relevance of this peak shift remain unclear.

### 4.2 Functional connectivity and signal variability

By charting patterns of connectivity across brain regions, FC analyses offer a detailed picture of how neural dynamics are altered in psychedelic states. Our primary finding of decreased (directed) information flow together with unchanged or increased (undirected) correlation speaks to a disintegration of communication between and within brain regions, which in turn implies a loosening of dynamical constraints on brain activity in the psychedelic state. This loosening may correspond to an increased repertoire of dynamical states, in line with theoretical proposals that link increased dynamical diversity to the characteristic subjective effects of psychedelics including unconstrained cognition, perception, and ego-dissolution (Carhart-Harris et al., 2014; Atasoy et al., 2018; Carhart-Harris, 2018; Carhart-Harris and Friston, 2019).

Our results are most directly comparable to other studies of the psychedelic state using EEG or MEG. In particular, in a previous analysis of the same dataset (except the tiagabine control), we found increased Lempel-Ziv complexity for all psychedelics (compared with placebo), which can be considered as a measure of signal diversity across time and/or space (Schartner et al., 2017). This increase in diversity is in line with an increased repertoire of brain dynamics during the psychedelic state. However, unlike the present study, this previous study did not analyse connectivity of any kind. Other EEG and MEG studies (Kometer et al., 2015; Alonso et al., 2015; Rivolta et al., 2015; Pallavicini et al., 2019) have applied FC measures, but did not systematically compare directed and undirected measures, and did not compare results across a range of psychedelics (see Introduction).

fMRI studies of altered FC and signal variability in the psychedelic state are more common than M/EEG studies. These fMRI studies have revealed a wide range of effects across a range of different psychedelics. Some reported effects support an increased dynamical repertoire in the psychedelic state. Among these studies are those which employ measures of signal variability. In a relatively early study, Tagliazucchi et al. (2014) report a wider repertoire of connectivity states following psilocybin administration. Lebedev et al. (2016) calculate voxel-level sample entropy from resting-state fMRI data, comparing LSD with placebo, and finding increased entropy in sensory and higher networks across multiple time scales. Viol et al. (2017) estimate functional brain networks from resting-state fMRI data recorded under ayahuasca, finding an increase in the entropy of the network degree distribution (a measure of network complexity) in the drug state. Atasoy et al. (2017) perform a “connectome-harmonic decomposition” of brain activity based on fMRI recordings under placebo and LSD (connectome harmonics are identified as spatial patterns reflecting synchronisation of activity at different spatial scales). They too report an expanded repertoire of active brain states under LSD. Collectively, these studies point to increases in the repertoire of neural dynamics during the psychedelic state, which is consistent with our finding of decreased FC in these states.

Other fMRI studies have focused on measures of undirected FC. These studies have revealed a range of altered patterns in the psychedelic state, including decreased coupling between ‘connectivity hub’ brain regions (Carhart-Harris et al., 2012); changes in connectivity within and across resting state networks that correlate with subjective effects (Carhart-Harris et al., 2016), and global increases in (undirected) FC in high-level association cortices and the thalamus, correlating with subjective reports of ego dissolution (Tagliazucchi et al., 2016). A number of fMRI studies have examined modifications of connectivity at sub-anesthetic doses of ketamine. However, a confusing pattern emerges in these papers, with some showing increases in connectivity (Driesen et al., 2013; Anticevic et al., 2015; Dandash et al., 2015), whereas others show decreases (Kraguljac et al., 2017; Wong et al., 2016). This pattern may be related to the many different analytical techniques to both preprocess and quantify FC in the BOLD data. These observations of increases and decreases in FC are difficult to directly compare with the present results, given the slow temporal dynamics of the fMRI BOLD signal, but they are by-and-large consistent with the idea of an increased repertoire of brain dynamics during the psychedelic state. We note that directed measures of FC, such as Granger causality, remain controversial when applied to fMRI because of the slow nature of the BOLD signal (Seth et al., 2013; Solo, 2016; Barnett and Seth, 2017).

### 4.3 Directed and undirected functional connectivity

A unique feature of the present analysis is the comparison of directed (Granger causality) and undirected (correlation and coherence) on the same data. This comparison is permitted by the high-temporal resolution, accurate source-localisation, and steady-state nature of the MEG recordings, which are ideally suited for the application of rigorous and robust directed FC analyses.

As already mentioned, our primary finding of decreased directed FC across all psychedelics (but not tiagabine) is in line with the notion of an increased repertoire of brain dynamics in the psychedelic state. There is a broader relevance to this finding, highlighted by the independent movement of directed FC and undirected FC (correlation and coherence) in our analyses. Had we examined only the more familiar measures of undirected FC, we would have drawn very different conclusions about the influence of psychedelics on global brain dynamics. In principle, directed FC measures (such as Granger causality) and undirected measures (such as correlation and coherence) offer distinct and independent perspectives on neural dynamics. Our study shows that this “in principle” difference also matters in practice. Future EEG/MEG studies of functional connectivity, both in the context of psychedelics and beyond, should therefore consider applying both directed and undirected FC measures in order to gain a comprehensive picture of neural dynamics, especially for exploratory analyses. It bears emphasising that FC analyses, whether directed or undirected, are distinct from EC analyses (EC; Friston et al., 2013; Seth et al., 2015). At the most general level, FC describes statistical dependencies between variables, whereas EC aims to identify the minimal causal circuit underlying some observed activity pattern (Barrett and Barnett, 2013). Intuitively, FC provides a description of dynamics, while EC [as operationalised by techniques such as dynamic causal modelling; see Friston et al. (2003)] provides an inference about underlying mechanism. Although the FC statistics we employ, such as Granger causality, are frequently operationalised via parametric modelling, these stochastic process models do not stand as mechanistic descriptions of neural dynamics, but as generic time-series models (Barnett et al., 2017). In FC approaches (both directed and undirected) the metrics derived have information-theoretic interpretations (Barnett et al., 2009; Barnett and Bossomaier, 2013), and since the generic models are comparatively low-dimensional, these approaches are well suited to exploratory analysis of the type performed in this study.

### 4.4 Limitations

Our analysis has several limitations. First, it is difficult to completely exclude that artefacts due to fine muscle movements may have affected the MEG signal differently in drug as compared with placebo conditions. This concern applies in particular to LSD, where it applies primarily to higher frequency bands - notably the *γ*-band. While caution should be therefore be applied to results in these high frequency bands, we note that our main findings apply across all frequency bands, providing reassurance that muscle artefacts cannot readily explain our data.

A second limitation, which again applies primarily to LSD, is that the MEG recordings represent only a brief snapshot of an eight-hour psychedelic experience. Further research is needed to examine any possible time-varying dynamics of information flow across an entire “trip”.

Thirdly, we note that tiagabine recordings (drug and placebo) were carried out with eyes closed, whereas the other recordings were carried out with eyes open. However, comparisons between drug and placebo did not mix eyes-open and eyes-closed conditions, reassuring that this difference is unlikely to have affected our conclusions. Finally, inherent limitations on statistical power due to comparatively small sample sizes may have prevented us characterising more fine-grained alterations in information flow engendered by psychedelics.

## 5 Summary

We measured directed and undirected functional connectivity in source-localised MEG data recorded while healthy human volunteers experienced a psychedelic state (LSD, psilocybin, or low-dose ketamine), placebo, or a control state (tiagabine). We found that the psychedelic state is associated with a general decrease in directed functional connectivity (information flow), sometimes (in particular for LSD) accompanied by an increase in undirected functional connectivity (correlation). These changes in directed functional connectivity were not explainable by accompanying changes in spectral power, signal-to-noise ratio, or other features of the underlying statistical models (residuals correlation). The generalised decrease in information flow we observed is consistent with notions of an increased dynamical flexibility or repertoire in the psychedelic state (Carhart-Harris et al., 2016; Carhart-Harris, 2018; Carhart-Harris and Friston, 2019).

The distinct and in some cases (LSD) opposite movement of directed and undirected measures suggests that analyses of functional connectivity should, especially when exploratory, analyse both kinds of measure in order to deliver a comprehensive picture of underlying neural dynamics. Future research would usefully probe how directed and undirected FC measures behave with respect to each other in other contexts, besides psychedelics. The methods described here provide a viable, statistically-sound pipeline for implementing such analyses given epoched, multivariate neurophysiological data with reasonable temporal and spatial resolution.

## Acknowledgements

AKS and LCB are grateful to the Dr. Mortimer and Theresa Sackler Foundation, which supports the Sackler Centre for Consciousness Science. AKS is additionally grateful to the Canadian Institute for Advanced Research (Azrieli Programme on Brain, Mind, and Consciousness). SDM is supported by a Rutherford Discovery Fellowship. RLCH is supported by the Alex Mosley Charitable Trust, Ad Astra Trust, Tim Ferriss, Singhal Health Foundation, and the Tamas family.

## A Correlation and Granger causality (GC) for a minimal VAR model

We start by examining the simplest bivariate system:

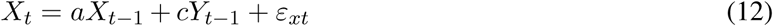

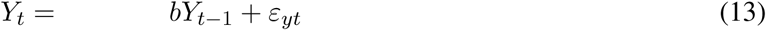

or

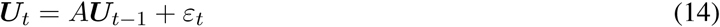

where 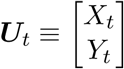 is the bivariate vector process, 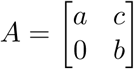 the VAR coefficients matrix and 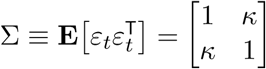 the residuals covariance matrix. The AR operator and MA operator (transfer function) are, respectively

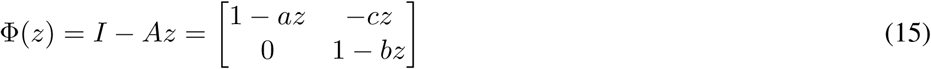

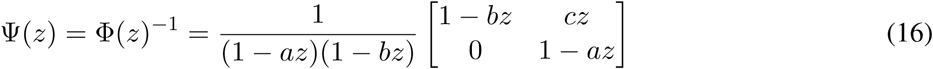

In the time domain, *z* may be interpreted as the backshift (lag) operator, while in the frequency domain *z* = *e*^*−iω*^ lies on the unit circle in complex plane, where *ω* is the phase angle, measured in radians. The CPSD may be calculated from the spectral factorisation (Wilson, 1972) *S*(*z*) = Ψ(*z*)∑Ψ(*z*)^*^ as

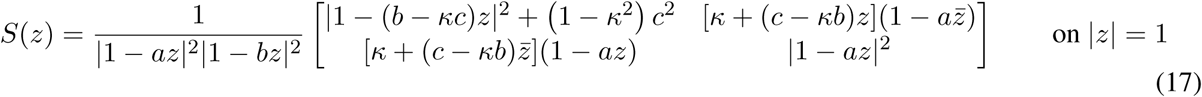

(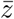 denotes complex conjugate) so that, in particular, the power spectral density of the sub-process *X*_*t*_ is

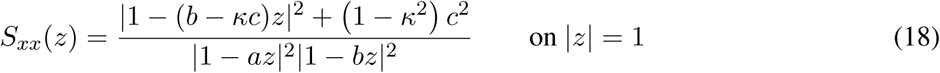

We factor this as

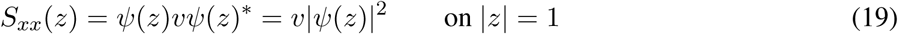

where 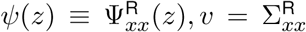 are, respectively, the MA operator and residuals variance for the reduced linear representation of the sub-process *X*_*t*_. By inspection, we try a reduced spectral factorisation with

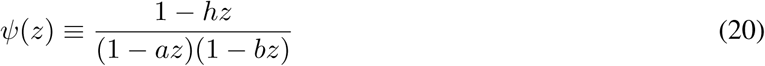

which yields

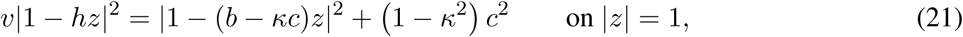

or

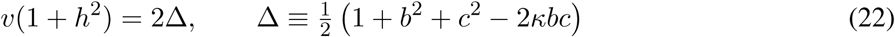

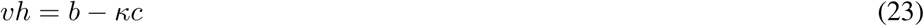

This yields the quadratic equation

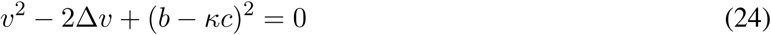

for *v*, so that

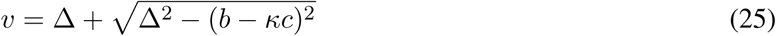

Note that we need the ‘+’ sign on the square root, since this yields correctly *v* = 1 when *c* = 0.

In the time domain, the mutual information between *X*_*t*_ and *Y*_*t*_ is given by

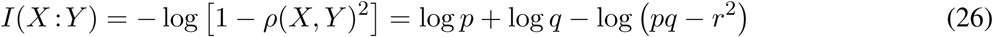

where *ρ*(*X, Y*) is the correlation between *X*_*t*_ and *Y*_*t*_, and 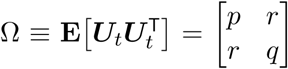 is the covariance matrix.To calculate Ω, we need to solve the discrete-time Lyapunov equation (derived from the Yule-Walker equations)

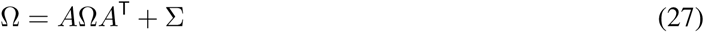

We find

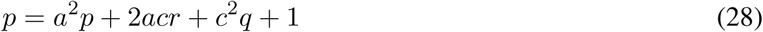

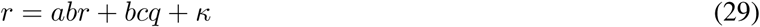

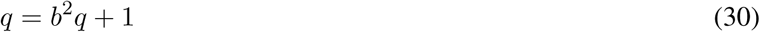

so

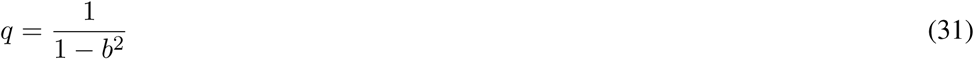

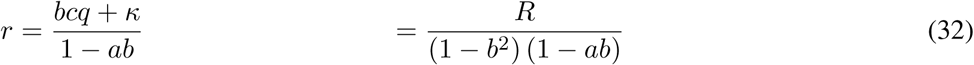

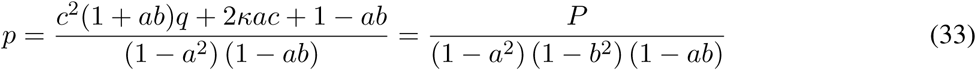

where

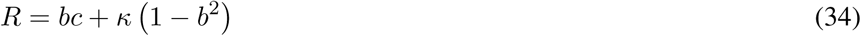

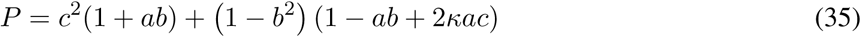

We then have

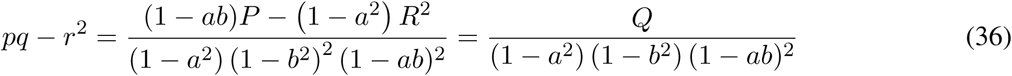

where

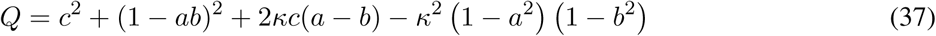

The MI is thus

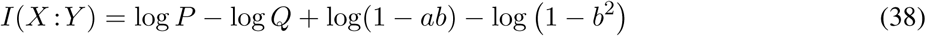

For the Granger causality from *Y* → *X*, since ∑_*xx*_ = 1, we have simply

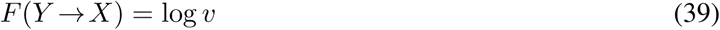

(the GC from *X* → *Y* is trivially zero). We note that the Geweke “total linear dependence” measure (Geweke, 1982, 1984) is given by

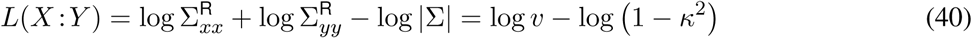

(note that 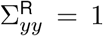). This also follows directly from Geweke’s decomposition of total linear independence into directional and instantaneous terms. We see immediately that if *κ* = 0 (zero residuals correlation), then *F* (*Y* →*X*) = *L*(*X* : *Y*).

In the frequency domain, we have, from the definitions,

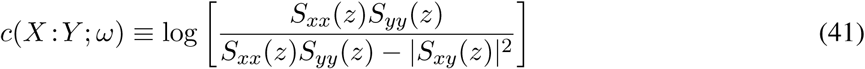

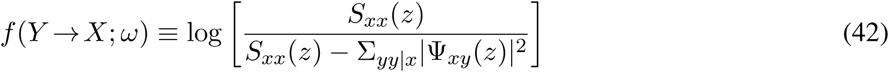

and straightforward calculation leads to

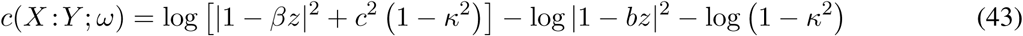

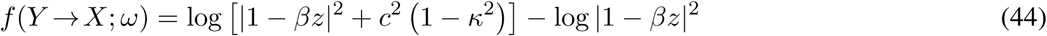

where *β ≡ b* − *κc*. We see immediately that if *κ* = 0 then *f* (*Y* → *X*; *ω*) = *c*(*X* : *Y* ; *ω*) at any frequency *ω*. Note that *f* (*Y* → *X*; *ω*) integrates to *F* (*Y* → *X*), while *c*(*X* : *Y* ; *ω*) integrates to *L*(*X* : *Y*).

We are, in particular, interested in regions and directions in parameter (*a, b, c, κ*)-space along which *I*(*X* : *Y*) and *F* (*Y* →*X*) move in opposite directions. To this end we may calculate the angle *θ* between the gradient vectors ∇*I*(*X* : *Y*) and ∇*F* (*Y* →*X*):

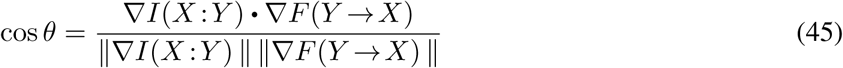

at a point (*a, b, c, κ*), where “•” denotes vector dot-product. The cosine is then +1 when the gradients are in exactly the same direction, −1 when they point in opposite directions and 0 when they are orthogonal. Gradients in the *κ* direction are of particular interest. Noting that 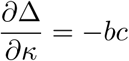, we may calculate:

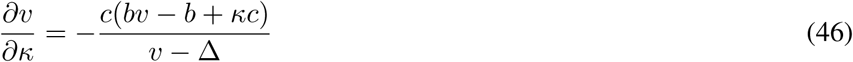

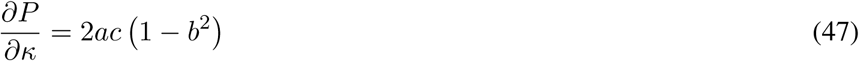

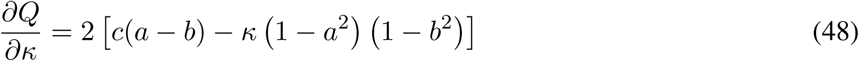

so that

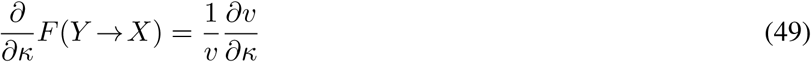

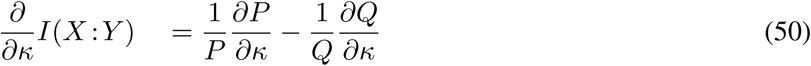

Numerically, we may establish that (except at *c* = 0) 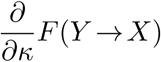 and 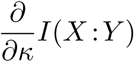 *always have opposite sign*; so, keeping the parameters (*a, b, c*) fixed, either increasing or decreasing *κ* always moves *I*(*X* : *Y*) and *F* (*Y* →*X*) in opposite directions. We stress, however, that we have not established that this necessarily applies in a highly multivariate scenario.

## Footnotes

1 We assume frequencies above 150 Hz are not biophysically relevant. In addition, downsampling improves some functional connectivity measurements (Barnett and Seth, 2017).

2 As a cross-check, we also independently estimated the undirected measures (partial correlation/coherence) from standard maximum-likelihood variance-covariance matrix estimates, and the cross-power spectral density (CPSD) matrix, estimated in sample by a multi-taper method with aggregation over epochs (Thomson, 1982; Bokil et al., 2010). Results were consistent with the VAR results.

3 Unlike partial correlation, mutual information is not signed; in the multivariate case, this would not make sense in any case.

4 A subtlety is that reduced VAR model parameters must be calculated directly from the full model; separate full and reduced regressions are known to induce strong biases, particularly in the frequency domain (Chen et al., 2006; Barnett and Seth, 2014; Barnett et al., 2017). Here we used a hybrid technique involving just the full regression (1), with reduced model parameters calculated using the state-space method of Barnett and Seth (2015).

5 For partial coherence/correlation, the analogue of (9) does *not* hold in general, except in the unconditional case

6 Preliminary mathematical analysis, which we intend to publish as a separate study, indicates that in general a decrease in SNR is associated with a decrease in both GC and MI.

7 Preliminary investigations suggest that increased residuals correlation may potentially (but not conclusively) indicate the presence of latent (unmeasured) common influences on the system dynamics; intuitively, linear modelling “sees” the effect of common—but unmodelled—inputs as correlated noise.

